# Glucose repression of *HXK1* is glucose flux-dependent via non-canonical regulation of Mig1

**DOI:** 10.64898/2026.08.09.743801

**Authors:** Ang Li, Michael Springer

## Abstract

Glucose is the preferred carbon source for budding yeast. Glucose sensing is achieved through multiple pathways, and the regulation of glucose-responsive genes has been reported to depend on both glucose concentration and glucose flux. However, the extent to which either of these mechanisms is used, and how cells sense glucose metabolic flux and couple it to transcriptional repression, remains unclear. Using tunable control of hexose transporters and hexokinases together with an optimized intracellular glucose sensor, we decoupled glucose uptake, phosphorylation, and intracellular glucose levels. We found that regulation of a Mig1-dependent reporter gene correlates with glucose flux rather than glucose concentration. Deletion of all known plasma membrane glucose sensors or replacement of yeast hexokinase with a bacterial glucokinase did not disrupt flux-correlated repression. Systematic mutational analysis of glucose signaling pathways showed that this Mig1-dependent response is mediated by the Snf1/AMPK pathway, but only at low glucose concentrations. At high glucose concentrations, Mig1 activity is controlled by an unknown, non-canonical mechanism. While consistent with much of the extensive literature on glucose regulation in *S. cerevisiae*, this work shows that careful quantitative analysis can uncover previously unrecognized modes of regulation.

## INTRODUCTION

Glucose is the preferred carbon source for the budding yeast *Saccharomyces cerevisiae*, and over evolutionary time, yeast has developed elaborate mechanisms to sense glucose and regulate its cellular activities in response to glucose. Extensive research has been conducted to dissect the molecular details of glucose sensing and regulation in yeast [1–5]. Glucose is sensed through both intracellular and extracellular mechanisms. Three plasma membrane-associated glucose sensors have been identified: the hexose transporter-like glucose sensors Snf3 and Rgt2 [6] and the G protein-coupled receptor (GPCR) Gpr1 [7, 8]. Several intracellular glucose-sensing systems have been described. The Ras2 protein responds to the glycolytic intermediate fructose-1,6-bisphosphate (FBP), indirectly coupling glucose concentrations to cAMP-protein kinase A (PKA) signaling [9]. Furthermore, glucose repression mediated by the transcriptional repressor Mig1 downregulates genes involved in the metabolism of alternative carbon sources [5, 10].

A central mystery in glucose signaling is that several glucose sensing pathways depend not only on glucose concentration, but also on glucose metabolic flux. For instance, recent studies have demonstrated that Ras2 senses FBP and could thereby link glycolytic flux to the PKA response [9]. However, the evidence does not strictly confirm a flux-based mechanism; rather, it shows that mutants with elevated glucose flux display increased FBP levels and elevated Ras2 activity. Similarly, activation of the Gpr1 signaling pathway requires glucose flux through hexokinase, as deletion of hexokinase abolishes signaling altogether [11]. Notably, the correlation between glucose repression and glucose phosphorylation activity has been noticed since 1989 [12], and flux-dependent regulation has subsequently been observed in a variety of cellular contexts [13, 14].

It is increasingly recognized that metabolic enzymes themselves can act as sensors [15]. For example, galactokinase functions as a flux sensor in the galactose utilization pathway [16]. Hexokinase, which catalyzes the first committed step of glycolysis, has long been postulated to serve a similar sensing function. Studies from the early 2000s hypothesized a role for hexokinase in glucose flux sensing [1, 17], and a signaling role for hexokinase has also been observed in higher plants [18]. Indeed, multiple attempts have been made to determine whether hexokinase itself acts as a key sensor in glucose repression [19]. Several mechanisms have been proposed, such as the formation of a Hxk2–Snf1–Mig1 complex [20], but these explanations have not achieved wide acceptance [21]. Empirical evidence challenging the hypothesis that hexokinase directly senses metabolic flux to control glucose repression has left the role of Hxk2 in glucose sensing unclear [22, 23].

Because of the complexity of glucose sensing and regulation in yeast and despite the considerable effort and progress in elucidating the functions of key factors such as Mig1, Snf1, and others, the exact mechanisms by which glucose metabolic flux is sensed and how these signals are integrated with glucose repression remain incompletely understood. Advances in high-throughput quantitative measurements, genome editing, and synthetic biology have made it possible to measure the quantitative behavior of complex pathways and to perform perturbation analyses to validate or refute hypotheses and better understand the system-level functions of genes and their emergent behaviors.

## RESULTS

### Transcriptional repression of *HXK1* gene quantitatively depends on glucose and requires *HXK2*

To quantitatively analyze the glucose response, we sought to create a transcriptional reporter capable of measuring glucose repression. Fluorescent reporters are ideal for this purpose because they enable high-throughput measurements using methods such as flow cytometry or microscopy, which are essential for building a systems-level understanding of signaling.

Historically, *SUC2* has been used as a reporter gene in yeast to monitor glucose-responsive regulation [24], because *SUC2* encodes a secreted invertase whose expression can be measured using a biochemical assay. To develop a fluorescent reporter of *SUC2*, we fused the *SUC2* promoter to the fast-maturing yellow fluorescent protein mVenus (the mVenus NB variant) [25]. We then integrated this reporter cassette into the genome, thereby minimizing variation in expression that could arise from differences in plasmid copy number.

Glucose repression, also known as catabolite repression [26], exhibits complex temporal dynamics. Over the course of minutes, multiple physiological changes occur [27, 28], whereas over several hours, the system establishes a new steady state [27, 29]. Our interest was in the steady-state response to glucose. Thus, we grew cells in YPD with 2% glucose, transferred them to minimal medium with 128 mM (2.3%) glucose for a further 16 h of outgrowth, washed the cells, and finally transferred them to media containing glucose concentrations ranging from 1 mM (0.02%) to 128 mM (2.3%). Reporter responses were then measured by microscopy after approximately 7 h (Figure 1A).

**Figure 1.**
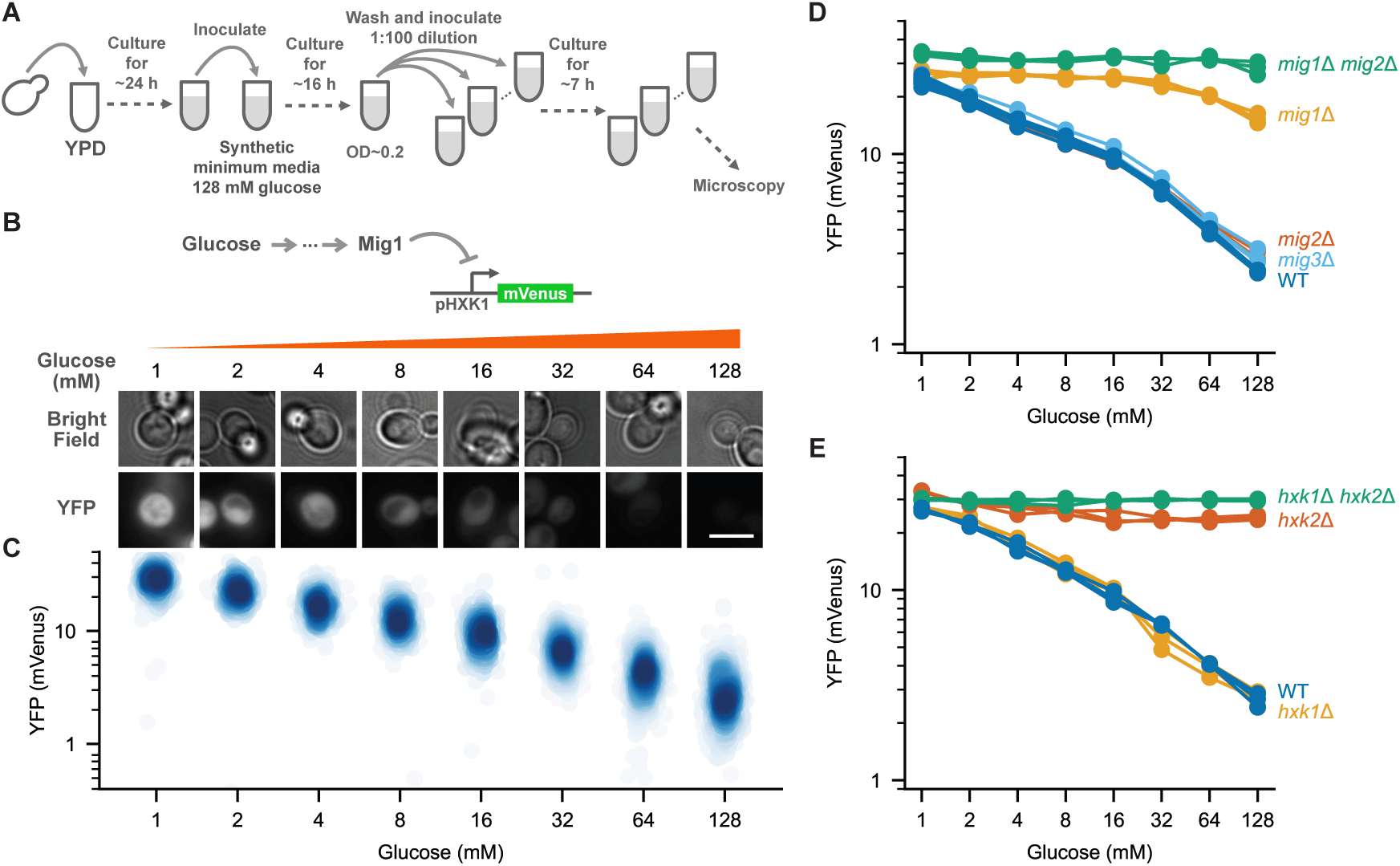
Quantitative measurement of glucose repression using an *HXK1* promoter fluorescent reporter. **(A)** Schematic of the steady-state glucose repression assay. Cells were grown to saturation in YPD (24 h), subcultured into synthetic minimal medium containing 128 mM glucose and grown to early logarithmic phase (16 h), washed, diluted 1:100 into synthetic minimal medium containing the indicated glucose concentrations (1–128 mM), and cultured for 7 h before fluorescence microscopy. **(B)** Schematic of the pHXK1-mVenus transcriptional reporter and representative bright-field and YFP images of reporter cells cultured at the indicated glucose concentrations. A pHXK1 promoter driving mVenus expression was used to quantify Mig1 transcriptional repression. Representative images with fluorescence intensities close to the population mean are shown. Quantification of fluorescence from the imaged cells is shown in **(C)**. Scale bar, 5 μm. **(C)** Single-cell YFP intensities of the pHXK1-mVenus reporter measured across the glucose titration. Each point represents one cell, *n* ≥ 356. X coordinates were randomly jittered for visualization. Three biological replicates were performed; data from one representative replicate are shown. Quantification of all three biological replicates is shown in **(E)** and labeled as WT. **(D)** Mean YFP fluorescence of the pHXK1-mVenus reporter in wild-type (WT), *mig1*Δ, *mig2*Δ, *mig3*Δ, and *mig1*Δ *mig2*Δ strains as a function of glucose concentration. Each line represents one biological replicate (*n* = 3). **(E)** Mean YFP fluorescence of the pHXK1-mVenus reporter in wild-type (WT), *hxk1*Δ, *hxk2*Δ, and *hxk1*Δ *hxk2*Δ strains as a function of glucose concentration. Each line represents one biological replicate (*n* = 3).

To quantify the response, cells were imaged using an epifluorescence microscope driven by custom high-throughput imaging software (Figure S2). Images were segmented using a custom U-net model optimized for yeast cell segmentation in bright-field images (Figure S3), and fluorescence intensities were quantified.

As expected, expression of the pSUC2-mVenus reporter responded to glucose (Figure S1B). However, despite the large fold change, the overall expression level from the *SUC2* promoter was low. As a result, the measurements of the fluorescent reporter were confounded by autofluorescence background. Such background limited the dynamic range of detection and the changes in the background could affect the reproducibility of results between measurements.

To mitigate the problems with the *SUC2* reporter, we aimed to develop a new, Mig1-dependent, transcriptional reporter for glucose repression that would be optimal for fluorescence-based measurements. Among genes reported to be regulated by Mig1, the hexokinase-encoding gene *HXK1* has a high expression level and is responsive to glucose repression [30]; other hexokinase isozymes have also been reported to be regulated by glucose repression and alternative carbon sources [31, 32]. To identify an improved glucose-responsive reporter, we tagged each of the three major hexokinase or glucokinase genes, *HXK1*, *HXK2*, and *GLK1*, with mScarlet-I at their native loci using a seamless gene-tagging method. Additionally, we tagged two hexokinase-like genes, *EMI2* and *NGK1* (YLR446W), which have some hexokinase activity [33, 34]. As a control for other factors that could affect a fluorescent reporter experiment, we expressed mScarlet-I under the constitutive *TDH3* promoter, which should not be regulated by glucose. We then cultured cells in media with varying extracellular glucose concentrations and measured steady-state reporter expression by microscopy with the same assay as in Figure 1A.

Based on microscopy analysis, *HXK1* exhibited the best dynamic range and signal-to-noise ratio. As expected, levels of Hxk1-mScarlet-I varied substantially across glucose concentrations (Figure S1A). Expression was low at 128 mM glucose but at least 10-fold higher at 1 mM glucose, with expression increasing monotonically as glucose decreased. Although *GLK1* and *EMI2* were also repressed by glucose, their fold changes were much smaller than that of *HXK1*. The remaining genes showed no significant glucose responsiveness (Figure S1A). We will refer to this glucose-dependent regulation of the *HXK1* promoter as glucose repression throughout the paper.

We then constructed a pHXK1-mVenus reporter by fusing the *HXK1* promoter to mVenus, integrated the construct into the yeast genome, and compared it with the pSUC2-mVenus reporter. When measured at steady state over a range of glucose concentrations, both reporters were regulated by glucose, but the pHXK1 reporter had a better signal-to-noise ratio than the pSUC2 reporter. The pHXK1 reporter responded to glucose repression in a graded, quantitative manner (Figure 1B), and the response was the same as that observed for the Hxk1-mScarlet-I protein fusion, indicating that *HXK1* expression is mainly regulated at the transcriptional level. We further performed a time course under our culture conditions and confirmed that the reporter was indeed measuring the steady-state response (Figure S1C and D).

To better understand the transcriptional regulation underlying glucose-dependent repression, we sought to confirm that *HXK1* regulation depended on *MIG1* and *HXK2*. We constructed *mig1*Δ and *hxk2*Δ mutants in our reporter strain and tested their glucose responsiveness. Deletion of either *MIG1* or *HXK2* largely abolished the response to glucose (Figure 1D and E). Repression between 128 mM and 1 mM glucose decreased from 10.1-fold in the wild-type strain to 1.7-fold in the *mig1*Δ strain and 1.4-fold in the *hxk2*Δ strain. To understand the remaining glucose response in the *mig1*Δ strain, we constructed a *mig1*Δ *mig2*Δ double-deletion strain because *MIG2* is known to contribute to repression in *mig1*Δ strains [30]. The *mig1*Δ *mig2*Δ double deletion strain completely abolished the responsiveness of the *HXK1* reporter over this concentration range (Figure 1D), resulting in only 1.2-fold repression between 128 mM and 1 mM glucose. Based on these results, we used the pHXK1-mVenus reporter for subsequent analyses throughout this study.

### Glucose repression of the *HXK1* gene correlates with glucose flux rather than glucose concentration

We wished to determine whether glucose repression depends on metabolic flux, glucose concentration, or a combination of the two. Several lines of evidence support a glucose flux-dependent mechanism: in *hxk2* mutant strains, hexokinase activity correlates with repression [12]; glucose transport is necessary for and correlates with glucose repression [35, 36]; and the glucose-6-phosphate isomerase reaction is required for glucose repression [37]. Conversely, other studies suggest that glucose repression is linked to either extracellular or intracellular glucose concentration rather than flux [38], with some proposing that free intracellular glucose is the signaling molecule [39]. Overall, distinguishing between flux- and concentration-dependent mechanisms is challenging because the two parameters are coupled, and both tend to affect cellular growth.

To address this question, we sought to create a strain in which we could independently control and manipulate glucose flux and intracellular glucose concentration, without altering the external glucose concentration. Modeling studies suggest that the glucose transport and hexokinase steps are the main points controlling glucose flux in glycolysis [40]. By perturbing these proteins—hexokinases for glucose phosphorylation and hexose transporters for glucose uptake—it should be possible to differentially affect glucose flux and intracellular glucose concentration [16]. Although both limiting either glucose uptake or glucose phosphorylation reduces glucose flux, these perturbations are expected to affect intracellular glucose concentrations differently. As predicted by metabolic modeling [40], restricting glucose uptake while maintaining hexokinase levels should lower intracellular glucose levels because less glucose is transported into the cell. In contrast, limiting glucose phosphorylation is expected to increase intracellular glucose concentration, provided that the rate of glucose import exceeds the capacity of hexokinase to phosphorylate glucose.

To control hexokinase activity, we deleted *HXK2*, which encodes the primary enzyme responsible for glucose phosphorylation [40]; this deletion alone is sufficient to abolish glucose repression (Figure 1E). Deletion of *HXK2* leads to increased expression of *HXK1*, which could confound our attempt to modulate glucose phosphorylation rates. To prevent changes in *HXK1* levels while titrating *HXK2*, we also deleted *HXK1* to eliminate this feedback. The *hxk1*Δ *hxk2*Δ double-deletion strain does not exhibit glucose regulation (Figure 1E) but is viable because glucose phosphorylation is provided by *GLK1* [41]. Unlike Hxk1, maximal glucose flux through Glk1 is limited by its polymerization mechanism [42]. We then constructed two strains in this *hxk1*Δ *hxk2*Δ background, with either *HXK1-*mScarlet-I or *HXK2-*mScarlet-I expressed from the doxycycline-inducible s promoter (Figure 2B).

**Figure 2.**
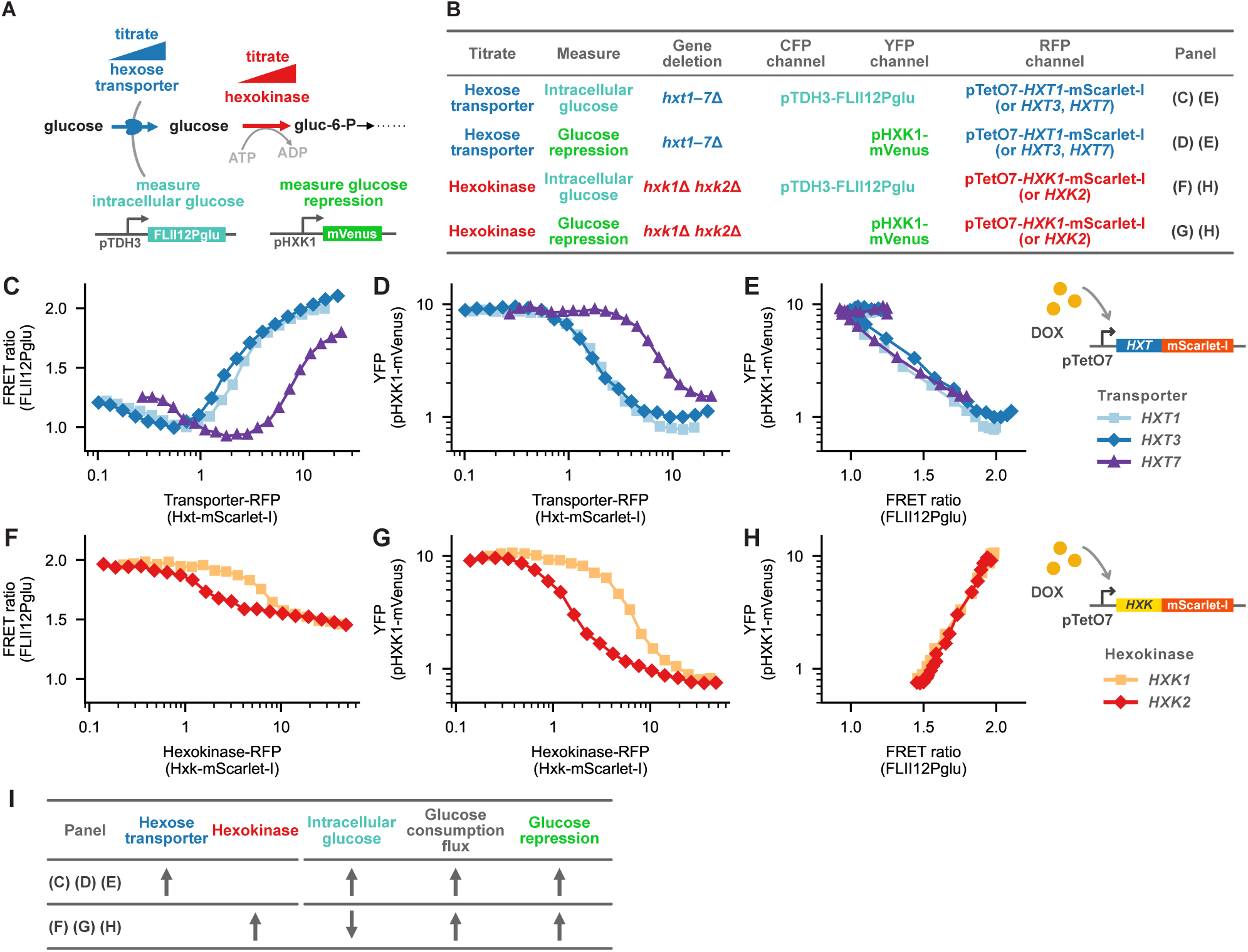
Glucose repression correlates with glucose flux rather than intracellular glucose concentration. **(A)** Experimental strategy used to independently perturb glucose transport and glucose phosphorylation. Hexose transporter expression was titrated to control glucose uptake, whereas hexokinase expression was titrated to control glucose phosphorylation. Intracellular glucose was measured using the FLII^12^Pglu FRET glucose sensor and glucose repression was measured using the pHXK1-mVenus transcriptional reporter. All cells were grown under glucose-rich conditions in 128 mM glucose. **(B)** Summary of the strains and fluorescent channels used for the transporter and hexokinase titration experiments. For transporter titration, *HXT1*–*HXT7* were deleted, and *HXT1*, *HXT3*, or *HXT7* was expressed from the doxycycline-inducible pTetO7 promoter as an mScarlet-I fusion. For hexokinase titration, *HXK1* and *HXK2* were deleted, and either *HXK1* or *HXK2* was expressed from pTetO7 as an mScarlet-I fusion. Intracellular glucose and glucose repression were measured in separate strains using pTDH3-FLII^12^Pglu and pHXK1-mVenus, respectively. **(C)** Intracellular glucose, measured as the FLII^12^Pglu FRET ratio, as a function of Hxt1-, Hxt3-, or Hxt7-mScarlet-I abundance during hexose transporter titration. Each point represents the average of at least 120 cells. **(D)** Glucose repression, measured as pHXK1-mVenus fluorescence, as a function of Hxt1-, Hxt3-, or Hxt7-mScarlet-I abundance during hexose transporter titration. Each point represents the average of at least 202 cells. **(E)** Relationship between glucose repression and intracellular glucose obtained by matching the transporter-expression titrations in **(C)** and **(D)**. **(F)** Intracellular glucose, measured as the FLII^12^Pglu FRET ratio, as a function of Hxk1- or Hxk2-mScarlet-I abundance during hexokinase titration. Each point represents the average of at least 198 cells. **(G)** Glucose repression, measured as pHXK1-mVenus fluorescence, as a function of Hxk1- or Hxk2-mScarlet-I abundance during hexokinase titration. Each point represents the average of at least 127 cells. **(H)** Relationship between glucose repression and intracellular glucose obtained by matching the hexokinase-expression titrations in **(F)** and **(G)**. **(I)** Summary of the transporter and hexokinase titrations and their effects on intracellular glucose concentration, glucose consumption flux, and glucose repression. Upward and downward arrows indicate increases and decreases, respectively.

To control glucose uptake, we deleted *HXT1*–*HXT7*, the collective set of hexose transporters responsible for the vast majority of glucose uptake [43]. These deletions also eliminate confounding effects on glucose uptake caused by feedback regulation of the other *HXT* genes. We then constructed three strains in this *hxt1–hxt7*Δ background with either *HXT1-*mScarlet-I, *HXT3-*mScarlet-I, or *HXT7-*mScarlet-I expressed from the doxycycline-inducible pTetO7 promoter.

To properly interpret our experiments, it was essential to directly measure intracellular glucose concentration. In the pTetO7-*HXK1* and pTetO7-*HXK2* strains, we titrated *HXK1 or HXK2* by varying doxycycline concentrations and confirmed by measuring glucose depletion that, as predicted, glucose flux was also titrated (Figure S4). To measure intracellular glucose concentrations, we constructed a strain expressing the FRET sensor FLII_12_Pglu [44–46]. This sensor is based on the *Escherichia coli* glucose/galactose-binding protein MglB and responds specifically to glucose and galactose [46]. Importantly, consistent with its crystal structure [47, 48], FLII_12_Pglu has been experimentally confirmed not to respond to glucose-6-phosphate [49].

Initial measurements with this glucose sensor suffered from a poor signal-to-noise ratio because of the low brightness of the sensor (Figure S5A–B). Because the sensor was not codon-optimized for yeast, we first attempted to improve it by replacing CFP and YFP with the codon-optimized and brighter CFP and YFP variants. However, these modifications reduced the FRET signal (Figure S5A–C). Ultimately, we codon-optimized the original ECFP-Citrine variant, resulting in a version with improved signal and signal-to-noise ratio (Figure S5A–C). To validate the glucose response of the sensor, we measured the FRET signal in digitonin-permeabilized cells across a glucose titration and showed that it correlated with glucose concentration (Figure S5D).

Finally, we used the pHXK1-mVenus reporter to measure glucose repression. Because of overlap between the fluorescent channels, we did not measure the glucose repression and the intracellular glucose concentration in the same cells during the glucose uptake or hexokinase titrations. Instead, we measured pHXK1-mVenus and FLII_12_Pglu(ECFP-Citrine) separately and matched the measurements based on glucose transporter or hexokinase levels. The strains we constructed enabled us to directly measure the relationship between intracellular glucose concentration and glucose repression while maintaining a constant extracellular glucose concentration (Figure 2A–B).

As predicted, increasing transporter levels led to higher intracellular glucose concentrations (Figure 2C). Similarly, increasing transporter expression increased glucose repression (Figure 2D). Titration of all three HXT transporters yielded the same underlying relationship between intracellular glucose and glucose repression (Figure 2E). The transporters displayed an offset in the relationship between transporter abundance and glucose repression (Figure 2D), which can be explained by the rate-affinity trade-off [36, 50]. Although precise values for *V*_max_, *K*_m_, and *k*_cat_ are not available for all three transporters, Hxt7 is known to be a high-affinity transporter, whereas Hxt1 and Hxt3 are low-affinity transporters. As a result, achieving the same intracellular glucose concentration requires approximately an order of magnitude higher level of Hxt7 expression than Hxt1 or Hxt3 expression.

As expected, increasing hexokinase expression reduced intracellular glucose concentrations (Figure 2F but still led to increased glucose repression (Figure 2G, Figure S6). Titration of both *HXK1* and *HXK2* produced nearly identical relationships between intracellular glucose and glucose repression (Figure 2H). The hexokinases also exhibited an offset in the relationship between their abundance and repression (Figure 2G). This offset can be explained by differences in their kinetic constants: Hxk2 has a higher *k*_cat_ than Hxk1 [40], so higher *HXK1* expression is required to match the effect of *HXK2* on intracellular glucose concentration.

The distinct relationships between intracellular glucose levels and glucose repression observed for transporter and hexokinase titrations (Figure 2E versus H) demonstrate that intracellular glucose alone cannot be the sole signaling molecule underlying glucose repression. This conclusion is strongly supported by the invariance of these relationships across titrations of the different transporters and hexokinases, respectively. Although the relationship holds over a wide range of transporter expression, we note that the decrease in intracellular glucose levels at very low transporter expression is unexpected. This result could be due to measurement limitations at low glucose concentrations or impaired cell growth at low transporter levels; however, it does not affect our main conclusion.

Although our data are inconsistent with intracellular glucose being the main regulator of glucose repression, they are consistent with regulation by glucose flux: perturbations predicted to increase or decrease glucose flux changed glucose repression in the same direction (Figure 2I). The data also remain consistent with other models, such as one in which a downstream molecule whose concentration correlates with glucose flux serves as the signal.

### Glucose sensors Snf3, Rgt2, and Gpr1 are not required for glucose repression of *HXK1*

Given that our data suggest that the *HXK1* promoter responds to a signal correlated with metabolic flux rather than intracellular glucose concentration, we set out to dissect the mechanism of this regulation. Specifically, we aimed to identify what was being sensed and which part of the complex glucose signaling network was responsible for transducing the signal. This is challenging because there are multiple glucose signaling pathways in yeast (Figure 3A), with cross-talk between branches and both intracellular and extracellular sensing systems (Figure 5A), making it difficult to determine how glucose is sensed.

**Figure 3.**
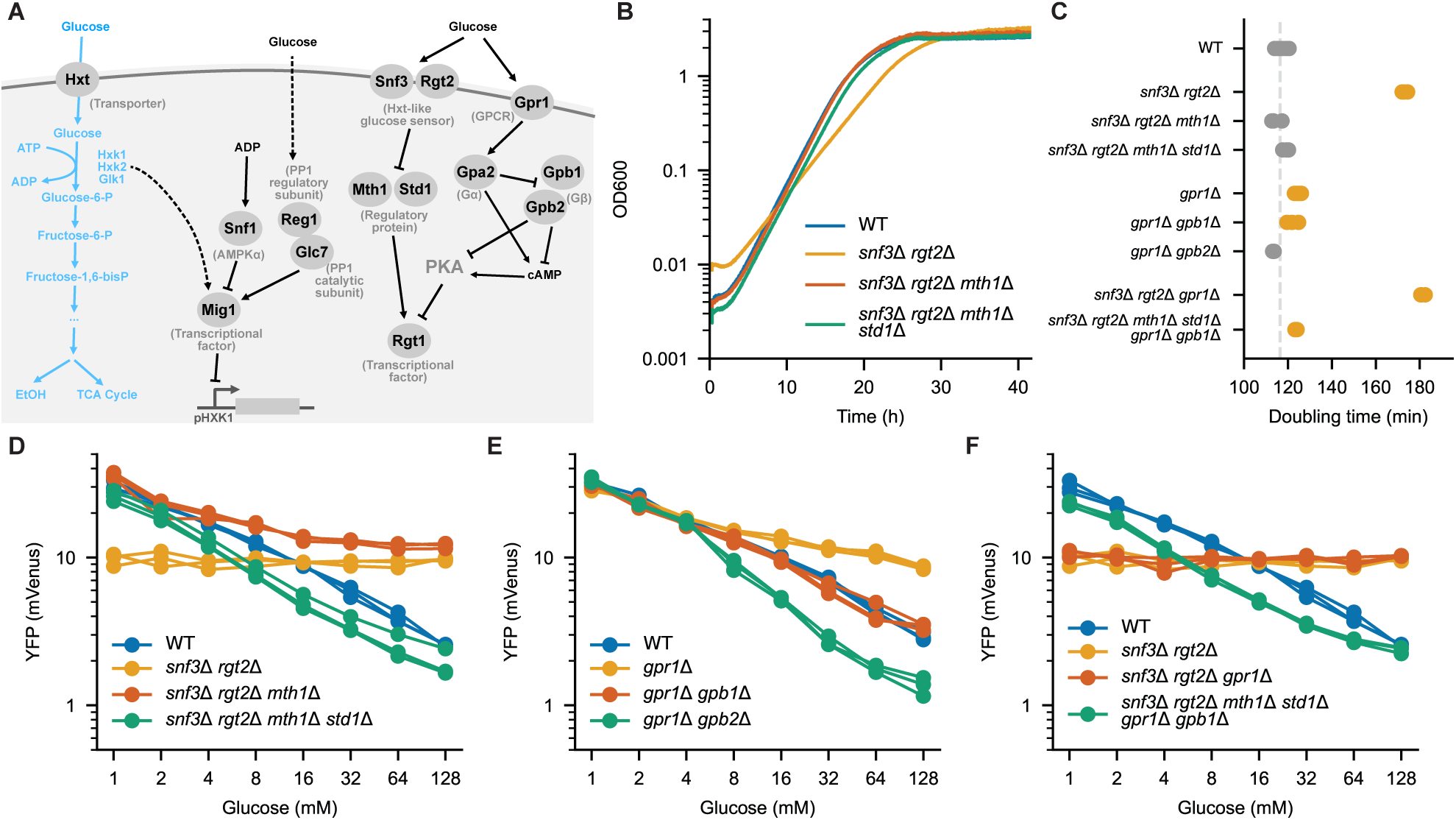
The extracellular glucose sensors Snf3, Rgt2, and Gpr1 are not required for glucose repression of the *HXK1* promoter. **(A)** Schematic of glucose metabolism and the major glucose signaling pathways. Blue labels indicate components involved in glucose transport and metabolism, whereas gray labels indicate known glucose signaling pathways that regulate transcription. The pHXK1-mVenus reporter was used to monitor Mig1-mediated glucose repression. **(B)** Representative OD_600_ growth curves of wild-type (WT) cells and strains carrying deletions in the Snf3/Rgt2 signaling pathway. OD_600_ was measured every 30 s using a custom growth measurement device. Three biological replicates were performed for each strain; one representative replicate is shown. Quantification of all biological replicates is shown in **(C)**. **(C)** Doubling times of strains carrying deletions in the Snf3/Rgt2 or Gpr1 signaling pathways. Each dot represents one biological replicate (*n* = 3). The dashed line indicates the doubling time of the wild-type strain. Orange points are significantly different from WT (*p* <0.001, independent two-sample *t* test). **(D)** Steady-state pHXK1-mVenus fluorescence as a function of glucose concentration in wild-type (WT), *snf3*Δ *rgt2*Δ, *snf3*Δ *rgt2*Δ *mth1*Δ, and *snf3*Δ *rgt2*Δ *mth1*Δ *std1*Δ strains. Each line represents one biological replicate (*n* = 3). **(E)** Steady-state pHXK1-mVenus fluorescence as a function of glucose concentration in wild-type (WT), *gpr1*Δ, *gpr1*Δ *gpb1*Δ, and *gpr1*Δ *gpb2*Δ strains. Each line represents one biological replicate (*n* = 3). **(F)** Steady-state pHXK1-mVenus fluorescence as a function of glucose concentration in wild-type (WT), *snf3*Δ *rgt2*Δ, *snf3*Δ *rgt2*Δ *gpr1*Δ, and *snf3*Δ *rgt2*Δ *mth1*Δ *std1*Δ *gpr1*Δ *gpb1*Δ strains. Each line represents one biological replicate (*n* = 3).

We hoped that by combining our glucose repression reporter with quantitative measurements, over a range of glucose concentrations, we could begin to unravel the mechanism. Although glucose repression correlates with flux, and we therefore expect it to depend on an intracellular signal, extracellular sensors are also known to influence expression of the *HXT* genes and, either directly or indirectly, glucose repression. Therefore, we first asked whether the extracellular glucose sensors directly contribute to the regulation of the *HXK1* promoter.

Our strategy was to remove glucose-sensing components singly and in combination, and then measure their effects on glucose repression of the *HXK1* promoter. A major complication is that mutations in the glucose signaling network can strongly affect growth, and growth defects or stress can themselves influence glucose repression. In the case of extracellular glucose sensors, deletion of these sensors converts cells to a low-glucose perceived state which can affect growth even when glucose levels are high [51, 52]. To address this, we verified mutant phenotypes by introducing compensatory mutations that restored cells to a high-glucose-perceived state without reestablishing signaling through the targeted pathway branch.

There are three extracellular glucose sensors in *Saccharomyces cerevisiae*: Snf3, Rgt2, and Gpr1. Snf3 and Rgt2 are homologs of glucose transporters and are thought to mainly control hexose transporter expression [6]. They achieve this through regulation of Rgt1 in concert with the cAMP/PKA signaling pathway [53]. To determine the potential role of these glucose sensors, we first created a pHXK1-YFP *snf3*Δ *rgt2*Δ strain. Using a custom growth measurement device that allows for highly accurate, continuous monitoring of cell growth, we found that this strain had a substantially decreased growth rate (Figure 3B and C). This is likely because these deletions cause the cells to perceive that they are in a low-glucose environment, even in glucose-rich media. Although this double-deletion strain completely abrogated glucose regulation, this could reflect either a direct or an indirect role of Snf3 and Rgt2 in the regulation of *HXK1* (Figure 3D). Indeed, glucose flux was significantly reduced in the *snf3*Δ *rgt2*Δ strain (Figure S7).

To distinguish between these two possibilities, we created two additional strains: one containing *mth1*Δ and the other containing both *mth1*Δ and *std1*Δ. Mth1 and Std1 act directly downstream of Snf3 and Rgt2 [54, 55]. They function as corepressors and transmit the glucose signals to the downstream transcription factor Rgt1 [56]. Deletion of *MTH1* and *STD1* should cause the cells to perceive a high-glucose state regardless of the presence or absence of Rgt2 and Snf3. Indeed, both mutants quantitatively restored the growth rate in the *snf3*Δ *rgt2*Δ strain background (Figure 3B and C). Compared with the pHXK1-YFP *snf3*Δ *rgt2*Δ strain, the *snf3*Δ *rgt2*Δ *mth1*Δ strain partially restored glucose repression of *HXK1*, whereas the *snf3*Δ *rgt2*Δ *mth1*Δ *std1*Δ strain completely restored glucose repression (Figure 3D). Together, these data rule out a direct role for Rgt2 and Snf3 in mediating the observed glucose dependence of the *HXK1* glucose repression.

Gpr1 is a plasma membrane G protein-coupled receptor (GPCR) that has been shown to sense glucose. Glucose binding leads to the activation of adenylate cyclase (Cyr1), which synthesizes cAMP and activates PKA. This leads to changes in cell size and regulation of filamentous growth, but it is unclear whether Gpr1 directly regulates *HXK1*. We created a pHXK1-YFP *gpr1*Δ strain and measured YFP expression over a range of glucose concentrations. Although the strain remained responsive to glucose, deletion of *GPR1* diminished the dynamic range of this response (Figure 3E). As with *SNF3* and *RGT2*, this effect could be indirect and caused by the changes in cell growth and therefore glucose flux, because this strain had a decreased growth rate (Figure 3C). To test this, we separately deleted each of the Gβ subunits, Gpb1 and Gpb2. Both the pHXK1-YFP *gpr1*Δ *gpb1*Δ and pHXK1-YFP *gpr1*Δ *gpb2*Δ strains largely recovered wild-type growth rates and the full range of glucose repression (Figure 3E). Thus, although Gpr1 responds to glucose, it is not required for the glucose-dependent repression of *HXK1*.

To fully rule out a role for the extracellular glucose sensors in regulating the graded glucose repression of the *HXK1* promoter we deleted all three glucose sensors together along with their downstream mediators, generating the pHXK1-YFP *snf3*Δ *rgt2*Δ *mth1*Δ *std1*Δ *gpr1*Δ *gpb1*Δ strain. This strain retained a similar glucose repression profile similar to that of the wild-type strain (Figure 3F) and had a growth rate near wild-type levels (Figure 3C). Taken together, these results indicate that the flux-correlated glucose repression response that we observed for *HXK1* expression is driven by an intracellular sensing mechanism.

### Hexokinase is not the direct glucose sensor for glucose repression

A range of phenotypic data supports the idea that the hexokinase Hxk2 contributes to glucose sensing. The connection between hexokinase activity and glucose repression was first reported in 1977 [57], and Hxk2 has since been identified as a key enzyme in this process [58]. In *hxk2* mutant strains, hexokinase activity correlates with the extent of glucose repression [12]. However, some studies have reported Hxk2 mutants in which signaling and catalytic activity appear to be decoupled [19, 59], suggesting that the enzyme’s signaling function may not be strictly tied to its catalytic role.

Despite extensive research, a molecular mechanism that definitively demonstrating the functions of Hxk2 in glucose signaling has not been established. Multiple molecular features of Hxk2 regulation have been described, including glucose-dependent nuclear localization [60, 61], carbon source-dependent phosphorylation at Ser14, and a dynamic monomer–dimer equilibrium [62–64]. However, it remains uncertain how these molecular phenomena relate to glucose sensing or signaling. Several mechanistic models have been proposed, such as one in which Hxk2 forms a complex with Snf1 and Mig1 [20], although this explanation is not universally accepted [21]. A simple mechanism has been uncovered in which the Gal1 galactokinase serves as a flux-correlated sensor, providing a previously untested mechanism that could also operate in glucose sensing [16].

Given that the mechanism of Hxk2 signaling remains unknown, we sought to directly test whether Hxk2 possesses a signaling function, independent of the assumed mechanism. For Hxk2 to act as a signaling factor, it would need to physically interact with a downstream signaling component. We reasoned that, unlike its interaction with glucose, which is highly conserved, specific interactions with signaling partners should not be conserved over long evolutionary distances because both molecules would have coevolved. Supporting this idea, replacing *S. cerevisiae GAL1* with *S. pombe GAL1* abolishes Gal1-dependent signaling without affecting its galactokinase activity [16].

To test whether Hxk2’s signaling function depends on such specific intermolecular interactions, we replaced *S. cerevisiae HXK2* with the *E. coli* glucokinase gene *glk* (EcGlk). Although most glucose in *E. coli* is metabolized through the phosphotransferase system, EcGlk encodes a glucokinase that catalyzes glucose phosphorylation. Critically, EcGlk phosphorylates glucose with kinetic parameters comparable to those of Hxk2 [65]. A previous study had replaced *HXK2* with human pancreatic glucokinase GlkB in *S. cerevisiae*, but neither glucose repression nor glucose-phosphorylation activity was restored to the same quantitative level, confounding interpretation [22]. Importantly, structural similarity between Hxk2 and EcGlk is restricted to the conserved catalytic core [66]. Their evolutionary distance and limited structural overlap make it highly unlikely that EcGlk would interact with the same downstream partners as Hxk2. Thus, replacing *HXK2* with EcGlk provides a stringent test of whether its potential signaling function depends on specific intermolecular interactions (Figure 4A).

**Figure 4.**
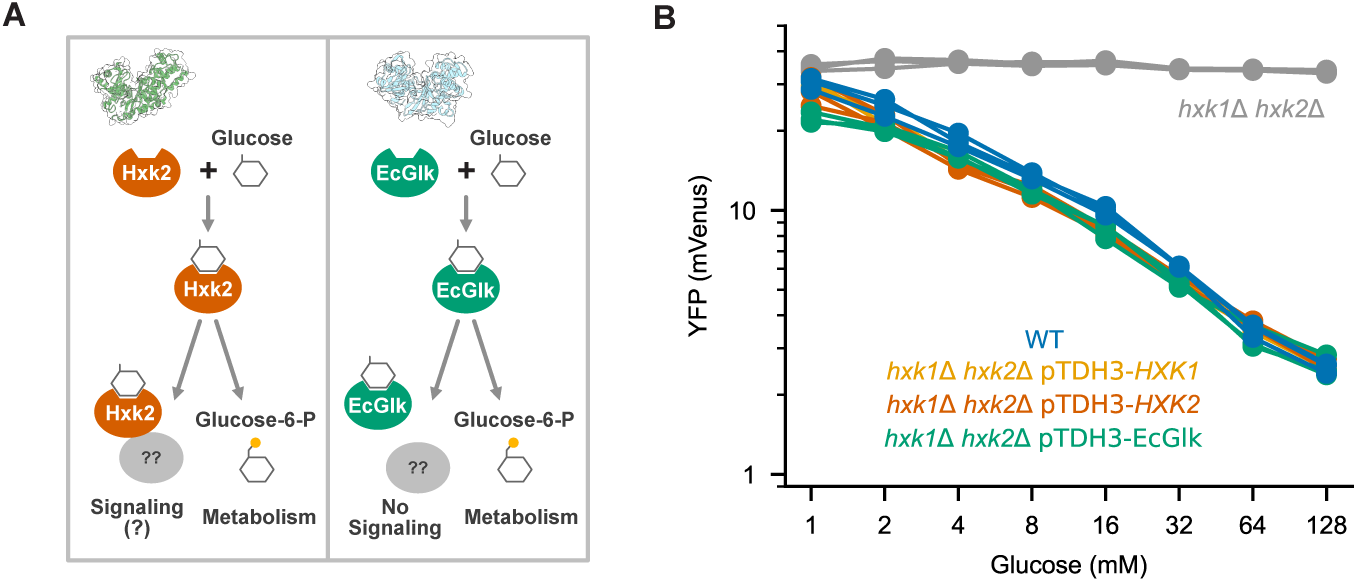
Hxk2-specific signaling is not required for flux-correlated glucose repression. **(A)** Experimental strategy for testing whether Hxk2 possesses a signaling function independent of its catalytic activity. Hxk2 has been proposed to directly couple glucose phosphorylation to glucose repression. To test this hypothesis, the endogenous yeast hexokinases were replaced with the evolutionarily distant *Escherichia coli* glucokinase EcGlk, which retains glucose-phosphorylation activity but, because of its evolutionary divergence, is unlikely to preserve specific protein–protein interactions with the yeast glucose-signaling network. **(B)** Steady-state pHXK1-mVenus fluorescence as a function of glucose concentration in wild-type (WT), *hxk1*Δ *hxk2*Δ, *hxk1*Δ *hxk2*Δ pTDH3-*HXK1*, *hxk1*Δ *hxk2*Δ pTDH3-*HXK2*, and *hxk1*Δ *hxk2*Δ pTDH3-EcGlk strains. Each line represents one biological replicate (*n* = 3).

We found that all three strains expressing either *S. cerevisiae HXK1*, *S. cerevisiae HXK2*, or *E. coli glk* from the strong *TDH3* promoter restored glucose repression to wild-type levels (Figure 4B). These results indicate that the proposed direct signaling function of Hxk2 is not required to explain glucose repression in yeast. Notably, although glucose repression was rescued in all strains, the growth defect was not rescued in the strain expressing EcGlk (Figure S8). This occurred even though glucose consumption flux in the EcGlk strain was comparable to that of the strains expressing *HXK1* or *HXK2*, which had growth rates close to that of the wild type (Figure S8). One possible explanation is that the yeast hexokinases are feedback-inhibited by trehalose-6-phosphate to limit the influx into glycolysis [67], whereas such inhibition may be absent for EcGlk.

Together, the data from Figure 2 and Figure 4 support the conclusion that flux-correlated repression of pHXK1 is regulated by a factor downstream of the hexokinases rather than by a hexokinase-intrinsic signaling function.

### No glucose regulatory pathway is solely responsible for the regulation of *HXK1*

Having ruled out both the plasma membrane glucose sensors and the hexokinases as sources of *HXK1* regulation, we next sought to determine whether other intracellular proteins or pathways known to participate in glucose signaling might account for the apparent flux-correlated regulation of *HXK1*. As outlined in the Introduction and schematized in Figure 5A, multiple signaling pathways contribute to glucose repression. To identify which, if any, were responsible for glucose-dependent repression of the *HXK1* promoter, we systematically deleted components of these pathways and measured the output of a pHXK1-YFP reporter across eight glucose concentrations. In total, we constructed and analyzed 47 single-deletion strains and 16 multi-deletion mutant strains.

**Figure 5.**
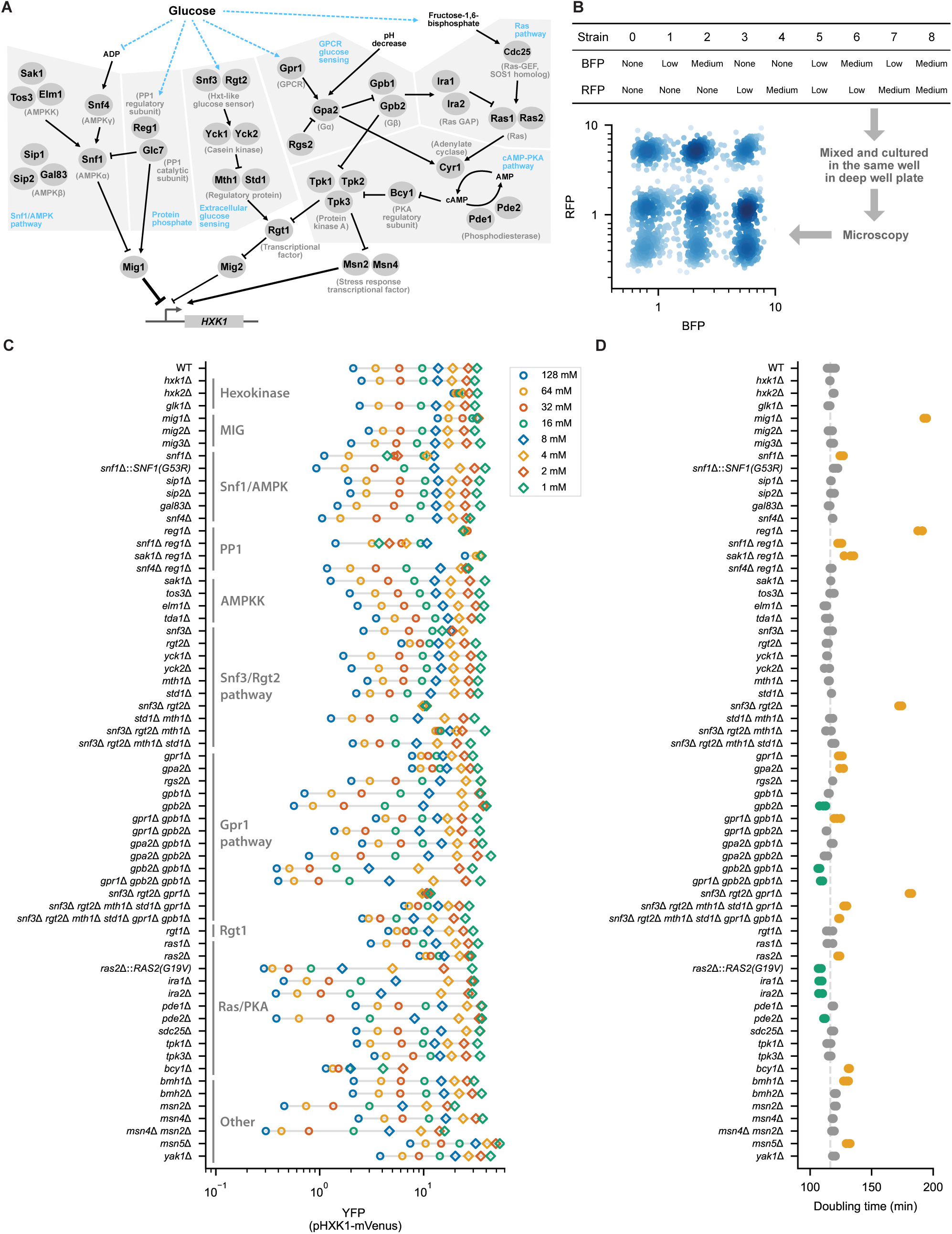
Systematic analysis of glucose signaling pathways regulating *HXK1* expression. **(A)** Schematic of the major glucose signaling pathways and regulators examined in this study, including the Snf1/AMPK, PP1/Reg1, Snf3/Rgt2, Rgt1, and Ras/cAMP/PKA pathways. Blue labels indicate pathways directly responsive to glucose or glucose metabolism. **(B)** Strategy for multiplexed fluorescence microscopy. Nine strains carrying unique combinations of BFP and RFP expression levels (none, low, or medium) were mixed and cultured in the same well. Following microscopy, individual strains were identified by their BFP and RFP fluorescence intensities. **(C)** Steady-state pHXK1-mVenus fluorescence measured across an eight-point glucose titration (1–128 mM) in wild-type and glucose signaling pathway mutants. Each point represents the mean fluorescence averaged across all biological replicates and independent isolates for the corresponding strain and glucose concentration. Three biological replicates were performed for each isolate, and at least two independent isolates were analyzed for each mutant. **(D)** Doubling times of wild-type and glucose signaling pathway mutants determined from OD_600_ growth curves. Each dot represents one biological replicate (*n* = 3 biological replicates per strain). The dashed line indicates the doubling time of the wild-type strain. Orange and green points indicate strains with significantly slower or faster growth than wild type, respectively (*p* < 0.001, independent two-sample *t* test).

To obtain high signal-to-noise data with internal standards while accommodating the large number of strains and conditions, we developed a multiplexed deep-well plate microscopy method that enabled simultaneous monitoring of nine strains per well. The nine strains in each well were distinguished by none, low, or medium expression of two fluorescent marker proteins, BFP and RFP (Figure 5B). These markers allowed unambiguous strain identification without measurably affecting growth. Eight wells were required to assay the nine strains across the eight glucose concentrations. Experiments were performed in triplicate using at least two independent isolates of each mutant. Measurements were performed under steady-state conditions, 7 h after transferring the cells to the target glucose concentration, with at least 1,800 cells analyzed per condition. To account for differences in growth rate among strains, more cells were added to the inoculation mixture when a mutant was expected to grow substantially more slowly. After demultiplexing, most strains had 100–400 cells analyzed per condition. Growth rates were measured in triplicate using custom high-precision growth devices.

Among all mutants tested, only *reg1*Δ eliminated glucose-dependent regulation. Reg1 forms a type 1 protein phosphatase (PP1) complex with Glc7 that dephosphorylates several key regulators of glucose signaling, including Snf1 [68], Mig1 [69], and Yck1 and Yck2 [70]. Similar to the *snf3*Δ *rgt2*Δ double mutant, deletion of *REG1* could have dominant effects on glucose regulation without necessarily being required for the core glucose-dependent repression mechanism. We therefore tested whether any secondary deletions could restore wild-type glucose regulation in the *reg1*Δ background, analogous to how *mth1*Δ *std1*Δ suppresses *snf3*Δ *rgt2*Δ, or *gpb1*Δ suppresses *gpr1*Δ. Indeed, we found that *snf4*Δ restored wild-type regulation of *HXK1* in a *reg1*Δ background. Thus, although the cAMP/PKA pathway, AMPK/Snf1 pathway, and Snf3/Rgt2 pathways can each exert dominant effects on glucose regulation, none of these pathways appears to mediate the core glucose-dependent repression of *HXK1*.

The interpretation that these three signaling pathways act as dominant tuners of a core glucose-sensing system regulating *HXK1* is further supported by the behavior of multiple single- and double-mutant strains. Several mutant strains altered the range or sensitivity of glucose repression. *snf1*Δ and *bcy1*Δ shifted the response range, remaining glucose repressible but showing uniformly lower *HXK1* promoter activity across all glucose concentrations. Conversely, *SNF1(G53R)*, *gpb1*Δ, *gpb2*Δ, *RAS2(G19V)*, *ira1*Δ, *ira2*Δ, and *pde2*Δ expanded the magnitude of repression, showing larger expression differences between 128 mM and 1 mM glucose than wild type. In contrast, *rgt2*Δ, *gpr1*Δ, *gpa2Δ*, *rgt1Δ*, and *ras2*Δ reduced the magnitude of repression, with smaller expression differences than wild type. Some mutants showed complex behaviors; for example, the *msn2*Δ *msn4*Δ double mutant both shifted and expanded the response range. Although these genes clearly modulate *HXK1* expression, none appears to be responsible for the core mechanism underlying glucose-dependent repression.

### Mig1 directly mediates flux-correlated glucose repression of the *HXK1* promoter

Having failed to identify any canonical upstream signaling pathway responsible for glucose sensing in the regulation of *HXK1*, we undertook a bottom-up approach and asked whether the glucose-dependent regulation of *HXK1* is directly mediated by Mig1. Although *HXK1* is known to be regulated by Mig1 [30], the evidence was limited to the observed derepression of *HXK1* in a *mig1*Δ strain. However, the *mig1*Δ strain exhibits a strong growth defect that complicates interpretation (Figure 5D). Thus, the loss of repression could be an indirect consequence of altered glucose metabolism.

We first performed mutational analysis of the *HXK1* promoter to determine whether the Mig1-binding sites are necessary for glucose-dependent regulation. The mutations were introduced into the pHXK1-YFP reporter construct, without affecting the regulation of endogenous *HXK1* or the growth of the strains. Conserved promoter motifs were identified by aligning orthologous sequences from *S. bayanus*, *S. kudriavzevii*, *S. mikatae*, and *S. paradoxus*, following established comparative genomic approaches [71]. Consensus binding sites for Mig1 and Msn2/4 were then mapped within conserved regions (Figure S9A). Combinatorial deletion of these sites confirmed that Mig1-binding sites 1 and 2 account for the majority of glucose-dependent repression, while Mig1-binding sites 3 and 4 also contribute to glucose-dependent repression and the Msn2/4 sites have little or no effect (Figure S9B and C). Deletion of all four Mig1-binding sites almost completely abolished glucose-dependent repression of the *HXK1* promoter (Figure 6A), reducing repression between 128 mM and 1 mM glucose from 11.5-fold for the wild-type promoter to 1.6-fold. These results demonstrate that the Mig1-binding sites are required for glucose-dependent regulation.

**Figure 6.**
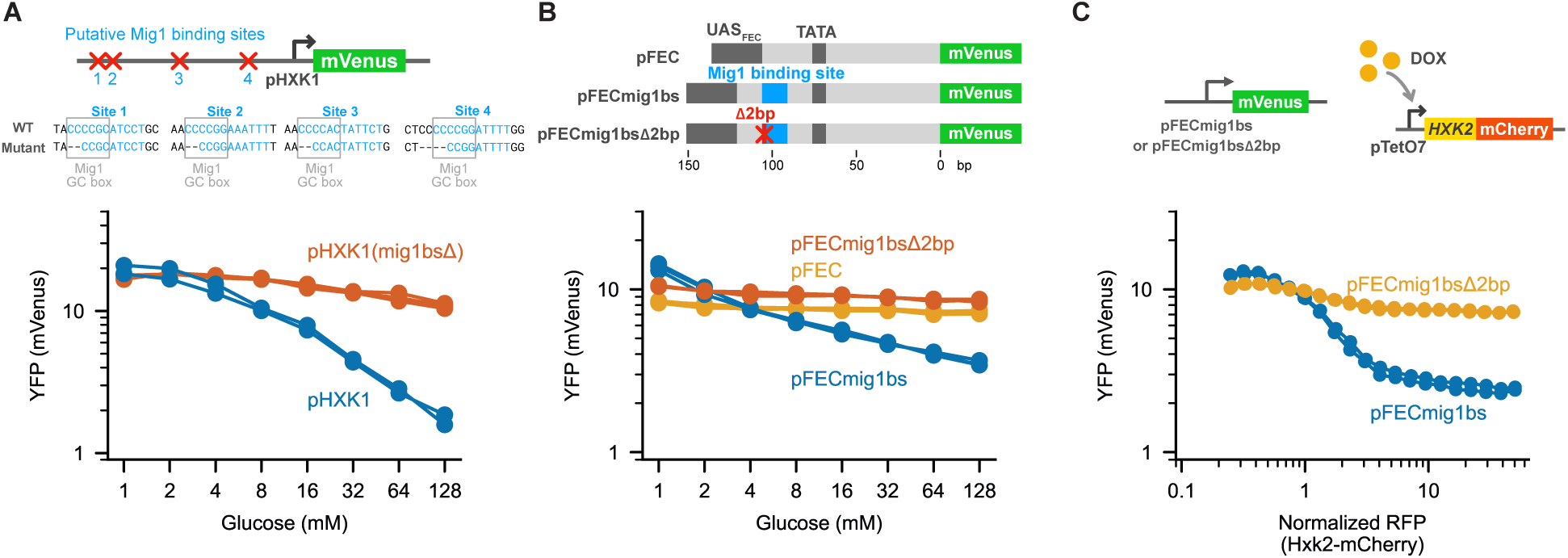
Mig1 directly mediates flux-correlated glucose repression of the *HXK1* promoter. **(A)** Mutation of putative Mig1-binding sites in the pHXK1-mVenus reporter. Top, locations of the four predicted Mig1-binding sites and the nucleotide deletions introduced into each site. Bottom, steady-state pHXK1-mVenus fluorescence as a function of glucose concentration for the wild-type promoter and the promoter carrying mutations in all four Mig1-binding sites (pHXK1(mig1bsΔ)). Each line represents one biological replicate (*n* = 3). **(B)** Design and characterization of a synthetic Mig1-responsive promoter. Top, schematic of the constitutive synthetic promoter pFEC, the Mig1-responsive derivative containing a single Mig1-binding site (pFECmig1bs), and a control promoter containing a disrupted Mig1-binding site with a 2 bp deletion in the GC box (pFECmig1bsΔ2bp). Bottom, steady-state fluorescence of the three reporters across a glucose titration. Each line represents one biological replicate (*n* = 3). **(C)** Response of the synthetic Mig1-responsive promoter to *HXK2* titration. In an *hxk1*Δ *hxk2*Δ background, *HXK2*-mCherry was expressed from the doxycycline-inducible pTetO7 promoter to vary Hxk2 level while cells were maintained in 128 mM glucose. Reporter fluorescence from pFECmig1bs or pFECmig1bsΔ2bp is plotted as a function of normalized RFP intensity. Each line represents one biological replicate (*n* = 2).

Next, to test whether a Mig1-binding site is sufficient to confer glucose regulation, we introduced a single Mig1-binding site into the fully synthetic promoter pFEC. This promoter contains synthetic upstream activating sequences (UAS_F_, UAS_E_, and UAS_C_), and exhibits constitutive expression [72]. As expected, the pFEC promoter was largely insensitive to glucose, exhibiting only 1.2-fold repression between 128 mM and 1 mM glucose (Figure 6B). Insertion of a single Mig1-binding site generated a robust glucose response, with the resulting pFECmig1bs reporter exhibiting 3.9-fold repression over the same glucose range. In contrast, introducing a two-base-pair deletion into the GC box of the Mig1-binding site (pFECmig1bsΔ2bp) reduced repression to 1.2-fold, abolishing glucose responsiveness. These results demonstrate that a Mig1-binding site is sufficient to confer glucose-dependent repression on a synthetic promoter.

Finally, to test the necessity and sufficiency of the Mig1-binding site in a flux-dependent context, we titrated *HXK2* expression and monitored reporter activity from the pFECmig1bs and pFECmig1bsΔ2bp constructs. Because varying *HXK2* levels modulates glucose flux under these conditions (Figure 2), we expected pFECmig1bs, but not pFECmig1bsΔ2bp, to respond to *HXK2* levels. To perform this experiment, *HXK2* was fused to mCherry and placed under the doxycycline-inducible pTetO7 promoter, enabling single-cell quantification of Hxk2 level and reporter activity. In cells grown in 128 mM glucose, the pFECmig1bs construct exhibited dose-dependent repression with increasing Hxk2 level (Figure 6C), with a dynamic range similar to that was observed with a glucose titration (Figure 6A). In contrast, pFECmig1bsΔ2bp showed little response to Hxk2 level.

Collectively, these results demonstrate that Mig1 directly mediates flux-correlated glucose repression of *HXK1*.

### Mig1 activity at high glucose is regulated by a non-canonical mechanism

The control of Mig1 transcriptional activity is generally thought to occur through modulation of its phosphorylation state and, consequently, its nucleocytoplasmic localization [3]. Under low-glucose conditions, Snf1/AMPK activity is high, leading to phosphorylation of Mig1 and its export to the cytoplasm by the exportin Msn5. As a result, Mig1-regulated genes are derepressed. In contrast, under high-glucose conditions, Snf1/AMPK activity is low, and Mig1 is dephosphorylated by the PP1/Reg1 complex. Consequently, Mig1 localizes to the nucleus and represses its target genes (Figure 7A).

**Figure 7.**
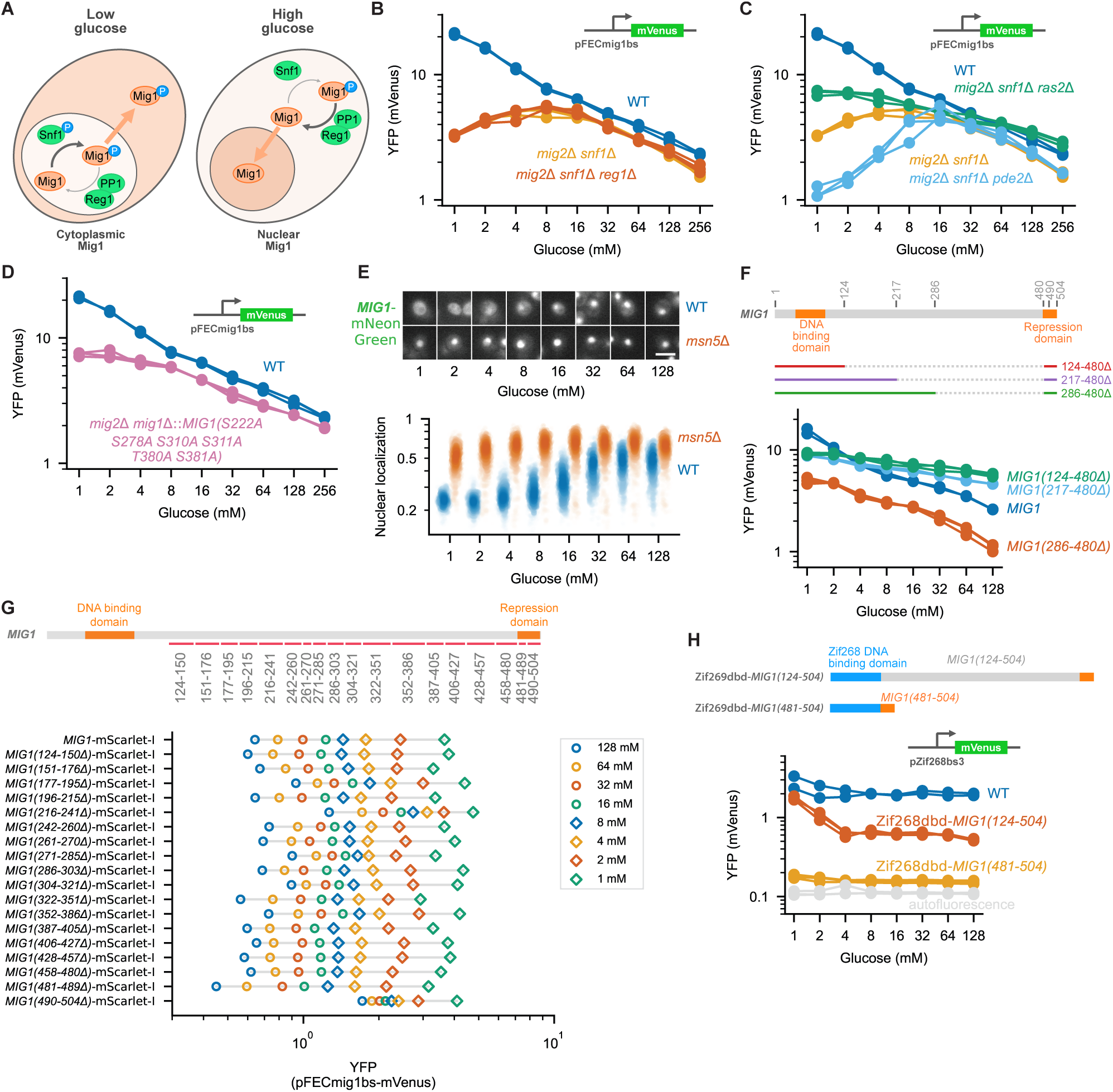
Mig1 activity at high glucose is regulated by a non-canonical mechanism. **(A)** Canonical model of Mig1 regulation. Under low-glucose conditions, Snf1/AMPK phosphorylates Mig1, promoting its export to the cytoplasm. Under high-glucose conditions, Mig1 is dephosphorylated by the PP1/Reg1 complex, accumulates in the nucleus, and represses transcription of its target genes. **(B-D)** Steady-state activity of the pFECmig1bs reporter across a glucose titration in **(B)** wild-type (WT), *mig2*Δ *snf1*Δ, and *mig2*Δ *snf1*Δ *reg1*Δ strains; **(C)** wild-type (WT), *mig2*Δ *snf1*Δ *ras2*Δ, *mig2*Δ *snf1*Δ, and *mig2*Δ *snf1*Δ *pde2Δ* strains; and **(D)** wild-type (WT) and a *mig2*Δ strain carrying the Mig1 phosphorylation-site mutant *MIG1(S222A S278A S310A S311A T380A S381A)*. Each line represents one biological replicate (*n* = 3). **(E)** Glucose-dependent localization of Mig1. Top, representative fluorescence images of endogenously tagged Mig1-mNeonGreen in wild-type (WT) and *msn5*Δ cells cultured at the indicated glucose concentrations. Bottom, single-cell quantification of Mig1 nuclear localization. Three biological replicates were performed; one representative replicate is shown. Scale bar, 5 μm. **(F)** Functional analysis of large Mig1 domain deletions. Top, schematic of the Mig1 domain-deletion constructs. Bottom, steady-state activity of the pFECmig1bs reporter in strains expressing the indicated Mig1 deletion mutants. Each line represents one biological replicate (*n* = 3). **(G)** Deletion scanning of Mig1. Top, schematic of the deletion-scanning strategy across the Mig1 regulatory region. Bottom, steady-state pFECmig1bs reporter activity in strains expressing individual Mig1 deletion mutants across the glucose titration. The mean of three biological replicates is shown. **(H)** Synthetic transcription factor analysis of Mig1 domains. Top, schematic of Zif268 DNA-binding domain fusion proteins containing either the Mig1 repression domain [Mig1(481–504)] or the complete regulatory region [Mig1(124–504)], assayed using the synthetic reporter pZif268bs3-mVenus. Bottom, steady-state reporter activity across the glucose titration. WT denotes cells lacking the synthetic transcription factor. WT data represent two biological replicates; strains expressing the synthetic transcription factors represent three biological replicates each.

To determine whether flux-dependent regulation of Mig1 activity arises from flux-dependent control of its phosphorylation state, we sought to characterize how Mig1 is regulated by upstream factors. We employed the pFECmig1bs reporter, which is specific for Mig1 activity. To simplify interpretation, we deleted *MIG2*, which encodes a Mig1 homolog that recognizes the same binding site. We then perturbed the upstream kinase Snf1/AMPK by deleting its catalytic subunit, *SNF1*.

Although deletion of *SNF1* altered Mig1-dependent regulation at low glucose concentrations (Figure 7B), the growth rate of this mutant was also reduced, as indicated by low cell counts (Figure S10), complicating interpretation. Surprisingly, *SNF1* deletion had little effect on glucose-dependent control of Mig1 activity at glucose concentrations above 16 mM (0.29%). The extent of repression between 256 mM and 16 mM glucose remained largely unchanged, increasing slightly from 2.7-fold in the wild-type strain to 3.0-fold in the *mig2*Δ *snf1*Δ strain. This finding suggests that Mig1 activity at high glucose is regulated by a Snf1/AMPK-independent, glucose-dependent signaling pathway.

To identify the origin of this glucose-dependent control of Mig1 activity in the *snf1*Δ strain, we first hypothesized that the signal might arise from the PP1/Reg1 complex, as glucose has been reported to activate PP1 phosphatase activity [73]. However, additional deletion of *REG1*, the regulatory subunit responsible for targeting PP1 to Mig1, in the *snf1*Δ strain failed to abolish the glucose response of Mig1 activity at high glucose concentrations (Figure 7B). Repression between 256 mM and 16 mM glucose remained 2.8-fold in the *mig2*Δ *snf1*Δ *reg1*Δ strain, comparable to the 2.7-fold repression observed in the wild-type strain. These results indicate that glucose signaling under these conditions is mediated by a pathway distinct from both Snf1/AMPK and PP1/Reg1.

One candidate pathway for the high glucose regulation of Mig1 at high glucose is the Ras/cAMP/PKA signaling cascade. Ras2 has been reported to couple glycolytic flux to PKA activation [9]. Although PKA is not known to directly phosphorylate Mig1, indirect regulation through other kinases or regulatory factors remains possible. If Mig1 were controlled by PKA, flux-dependent regulation of its activity could be readily explained. To test this hypothesis, we perturbed PKA activity by deleting either *RAS2* or *PDE2*. Deletion of *RAS2* reduces cAMP production and decreases PKA activity, whereas deletion of *PDE2* reduces cAMP degradation and increases PKA activity. However, neither *ras2*Δ nor *pde2*Δ abolished the control of Mig1 activity in the high-glucose range (Figure 7C). Repression between 256 mM and 16 mM glucose remained 3.0-fold in the *mig2*Δ *snf1*Δ *pde2*Δ strain, comparable to the 2.7-fold repression observed in the wild-type strain. Although repression was reduced to 1.8-fold in the *mig2*Δ *snf1*Δ *ras2*Δ strain, Mig1 activity remained glucose responsive.

Mig1 localization is regulated by phosphorylation at several sites, including S222, S278, S311, and S381. To determine whether the sites known to mediate Snf1-dependent regulation are important for the glucose-dependent regulation we observed, we created a Mig1 mutant lacking the Snf1 phosphorylation sites, Mig1(S222A S278A S310A S311A T380A S381A), in a *mig2*Δ background. This mutant was reported to be constitutively nuclear and to function as a constitutive repressor [74]. Again, we observed distinct behavior above and below 16 mM extracellular glucose. Below 16 mM, this mutant eliminated the glucose-dependent regulation, consistent with the roles of Snf1 and Reg1 below 16 mM glucose (Figure 7B and C). Surprisingly, rather than acting as a constitutive repressor, this Mig1 mutant remained glucose-responsive above 16 mM (Figure 7D). Repression between 256 mM and 16 mM glucose remained 2.4-fold, only modestly reduced from the 2.8-fold repression observed in the wild-type strain.

To determine whether the 16 mM glucose concentration corresponds to the concentration at which Snf1 has a physiological effect on Mig1 localization, we tagged the endogenous copy of *MIG1* with mNeonGreen at its C-terminus. Indeed, Mig1 localization transitioned around 16 mM glucose (Figure 7E). To confirm that this change reflected nucleocytoplasmic localization, we deleted the exportin Msn5, which is required for Mig1 nuclear export [74]. Consistent with previous reports, deletion of *MSN5* resulted in constitutive nuclear localization of Mig1 across the full glucose range (Figure 7E). Collectively, these mutant analyses strongly support the conclusion that glucose-dependent regulation of Mig1 at high glucose concentrations, above 16 mM, is mediated by an unknown, non-canonical mechanism.

To investigate how Mig1 is regulated by this potential non-canonical mechanism, we constructed a series of domain-deletion variants and synthetic transcription factor variants to identify regions required for glucose responsiveness. Mig1 is an intrinsically disordered protein. Previous studies established that the N-terminal region (residues 1–234) contains the zinc-finger DNA-binding domain, the internal region (residues 124–480) contains regulatory domains, and the C-terminal region (residues 481–504) contains the repression domain, including four critical hydrophobic residues [75–77].

First, we generated three internal domain deletion variants (286–480Δ, 217–480Δ, and 124–480Δ) with progressively larger deletions while preserving the N-terminal DNA-binding domain and the C-terminal repression domain (Figure 7F). In addition, we constructed a repression-domain deletion, 490–504Δ, which removes the critical hydrophobic residues in the C-terminal region. To assess the activity of these mutants, we used a pFECmig1bs reporter strain lacking *MIG2*, in which the endogenous *MIG1* locus was replaced with each mutant construct. Among the deletion variants, the 286–480Δ variant retained glucose responsiveness, exhibiting 2.5-fold repression between 128 mM and 16 mM glucose, although it was more strongly repressed than the wild-type strain, which exhibited 1.9-fold repression. In contrast, the 217–480Δ variant showed substantially reduced glucose responsiveness, with only 1.3-fold repression over the same glucose range (Figure 7F). These results suggest that residues 217–285 are required for glucose-dependent regulation of Mig1.

Next, we performed domain-deletion scanning across Mig1. Excluding the N-terminal DNA-binding domain (residues 1–123), we divided the remaining region (residues 124–504) into a series of small segments spanning 9–35 amino acids and generated mutants lacking each segment individually (Figure 7G). Mutant activity was measured using the pFECmig1bs reporter in a *mig2*Δ *mig3*Δ background in which endogenous *MIG1* was replaced with each deletion construct. Remarkably, most deletion mutants retained a normal glucose response in the high-glucose range. The only mutant exhibiting a strong defect lacked the C-terminal region spanning residues 490–504 (Figure 7G). Notably, none of the mutants spanning the 217–285 region (216–241Δ, 242–260Δ, 261–270Δ, and 271–285Δ) showed a substantial loss of glucose responsiveness at high glucose concentrations. This suggests that either the entire 217–285 region must be deleted to disrupt regulation or that this region functions together with residues 286–480.

Finally, we constructed synthetic transcription factors containing Mig1 domains to identify regions sufficient for glucose-responsive regulation. To test whether the C-terminal repression domain alone could confer glucose responsiveness, residues 481–504 of Mig1 were fused to the Zif268 zinc-finger DNA-binding domain (Zif268dbd), with residues 481–489 serving as a linker. The resulting fusion protein was expressed under the constitutive pSWI5 promoter. Activity was measured using a synthetic reporter containing three Zif268-binding sites (pZif268bs3). As expected, basal expression of the reporter was glucose-insensitive in the absence of the synthetic transcription factor (Figure 7H), exhibiting no detectable response in the high-glucose range, with a 1.0-fold change between 128 mM and 16 mM glucose, and only a modest response in the low-glucose range, with a 1.4-fold change between 8 mM and 1 mM glucose. The Zif268dbd–Mig1(481–504) fusion protein functioned as a repressor and reduced basal reporter expression, but likewise remained largely glucose insensitive, exhibiting 1.2-fold and 1.5-fold changes over the high- and low-glucose ranges, respectively. These results indicate that the C-terminal repression domain alone is insufficient to mediate glucose-responsive regulation.

To determine whether inclusion of the internal regulatory region could restore glucose responsiveness, we generated a Zif268dbd–Mig1(124–504) fusion protein, effectively replacing the native N-terminal DNA-binding domain of Mig1 with the heterologous Zif268 DNA-binding domain. This construct was assayed using the same reporter system. Inclusion of the internal domain partially restored glucose responsiveness, but only in the 1–4 mM glucose range (Figure 7H), which is likely related to the activation of Snf1/AMPK.

Together, these analyses indicate that the N-terminal DNA-binding domain (residues 1–123), the 217–285 region within the internal domain, and the C-terminal repression domain are all required for glucose-responsive regulation of Mig1 repression activity in the high-glucose range, whereas individual small segments within the internal region are largely dispensable.

## DISCUSSION

Glucose-dependent regulation in yeast has been extensively studied in a variety of contexts using diverse read-outs, including transcriptional responses, changes in protein localization, protein phosphorylation states, activation of signaling pathways such as kinases and phosphatases, intracellular pH changes, and metabolite dynamics. Previous genetic screens identified numerous genes and regulatory pathways involved in glucose signaling, as well as extensive interconnections among these pathways within the cellular signaling network [1–5]. However, it remains challenging to determine how individual signaling branches quantitatively contribute to the graded, glucose-dependent repression of Mig1-regulated genes. In this work, we built upon the extensive knowledge of the yeast glucose signaling network to investigate the quantitative behavior of Mig1-dependent glucose repression and the contributions of distinct regulatory pathways in this context.

Our results provide evidence that glucose repression of pHXK1 by Mig1 correlates with glucose flux rather than intracellular or extracellular glucose concentration. Although we were unable to directly measure glucose flux across the full range of glucose concentrations because of technical limitations associated with glucose consumption-based flux measurements, we were able to control flux by tuning the levels and identities of transporters or kinases. We confirmed the effects of these changes by measuring intracellular glucose while holding extracellular glucose constant. The opposing effects of hexose transporter and hexokinase titration on intracellular glucose, combined with predictable shifts in pHXK1 regulation among different hexose transporters and hexokinases, further supported the conclusion that the Mig1-mediated glucose response correlates with flux.

After eliminating a class of proposed models for glucose sensing, we further showed that the hexokinase Hxk2 is not essential for glucose flux sensing, as its function can be replaced by another enzyme with comparable glucose phosphorylation activity, such as *E. coli glk*. This finding suggests that flux-dependent regulation is mediated by a downstream metabolite or sensed at a downstream metabolic step as proposed in alternative models of glucose flux sensing [78, 79]. We did not attempt to identify the relevant metabolite or reaction in this study. Identification of the underlying sensing mechanism will be important for understanding the biochemical basis of flux-dependent regulation.

To determine how this flux-correlated regulation is mediated, we systematically examined known extracellular glucose sensors, major pathways upstream of Mig1 and Mig2, and additional genes and pathways previously implicated in glucose regulation by analyzing deletion mutants using the pHXK1 reporter. We ruled out regulation by known extracellular sensors by showing that a *snf3*Δ *rgt2*Δ *mth1*Δ *std1*Δ *gpr1*Δ *gpb1*Δ strain retained near-wild-type glucose-dependent regulation. We found that although the cAMP/PKA pathway and the Snf1/AMPK pathway can affect glucose regulation, neither appeared to mediate the core glucose-dependent repression of *HXK1*. These observations suggest either the existence of an additional major glucose-sensing pathway upstream of Mig1 or the possibility that Mig1 itself directly participates in glucose sensing.

To investigate the basis of this regulation, we measured Mig1 repression activity using the synthetic promoter pFECmig1bs and found that regulation of Mig1 phosphorylation and nucleocytoplasmic localization explains changes in Mig1 activity below, but not above, 16 mM extracellular glucose. These findings are consistent with the upstream signaling mutant analyses performed using the pHXK1 reporter. Because Mig1 activity could not be explained solely by localization changes induced by Snf1, an additional regulatory mechanism must contribute to the modulation of its repression activity. Domain analyses further showed that many small regions, and even substantial portions of the internal domain, are dispensable for glucose responsiveness. This observation is consistent with the intrinsically disordered nature of Mig1. At the same time, multiple domains were required simultaneously for proper regulation, suggesting that control of the Mig1 repression domain is not localized to a single regulatory motif or structured domain.

Although we were unable to determine the mechanism by which glucose regulates Mig1 above 16 mM extracellular glucose, several interesting possibilities remain. Previous studies reported that Mig1 can form oligomeric assemblies stabilized by interactions between intrinsically disordered regions [80, 81]. An earlier study using sucrose gradient sedimentation of whole-cell extracts identified a higher-molecular-weight form of Mig1 specifically in glucose-repressed cells [82]. One intriguing possibility is that a glycolytic intermediate could directly regulate Mig1 cluster or condensate formation. Because some glycolytic metabolites in yeast can accumulate to millimolar concentrations [40], they may be capable of inducing conformational or assembly-state changes in intrinsically disordered proteins such as Mig1, and these changes may drive cluster or condensate formation. Such a mechanism would connect previous phenomenological and modeling work on flux sensing to a potential mechanism through which this flux sensing regulates transcription. Investigation of this and other possible mechanisms may help explain how glucose flux regulates Mig1 repression activity.

## RESOURCE AVAILABILITY

### Lead contact

Requests for further information and resources should be directed to and will be fulfilled by the lead contact, Michael Springer.

### Materials availability

Plasmids and yeast strains generated in this work are available from the lead contact with a completed materials transfer agreement. Plasmid sequences are at Dryad in GenBank format at 10.5061/dryad.08kprr5j7.

### Data and code availability

All microscopy, growth curve data, data analysis code for reproducing figures in this study are deposited at Dryad at 10.5061/dryad.08kprr5j7 and are publicly available as of the date of publication.

The U-Net model finetuned for yeast cell image segmentation is also available at GitHub at https://github.com/angli339/unet_yeast_bf, and the weights of the model are available at Hugging Face at https://huggingface.co/angli339/unet_yeast_bf.

The high-throughput image acquisition software developed in this work is available open source at GitHub at https://github.com/angli339/NikonTiCtrl. The API library for controlling Tecan liquid handler in this work is available open source at GitHub at http://github.com/angli339/tecan-fluent-api.

Any additional information required to reanalyze the data reported in this paper is available from the lead contact upon request.

## ACKNOWLEDGMENTS

We thank the members of the Springer lab for discussions. This work was supported by a National Institutes of Health grant (R01 GM148497) to M.S.

## AUTHOR CONTRIBUTIONS

A.L. and M.S. conceptualized the study; A.L. developed the methods and performed the experiments; A.L. performed the data analysis and produced the figures; A.L. and M.S. wrote the paper; A.L. and M.S. reviewed and edited the paper; M.S. supervised the study.

## DECLARATION OF GENERATIVE AI AND AI-ASSISTED TECHNOLOGIES IN THE WRITING PROCESS

During the preparation of this work, the authors used ChatGPT in order to improve readability and language of the manuscript. After using the tool, the authors reviewed and edited the content for accuracy and take full responsibility for the content of the publication.

## SUPPLEMENTAL INFORMATION

- Document S1. Figures S1–S12
- Tables S1. Yeast strains used in this study
- Tables S2. Plasmids used in this study
- Tables S3. Key oligonucleotides used in this study

## EXPERIMENTAL MODEL AND STUDY PARTICIPANT DETAILS

### Bacterial strains and growth media

All plasmid construction steps were performed using *Escherichia coli* DH5α. For plasmid propagation, *E. coli* cells were grown in LB medium supplemented with the appropriate antibiotic (ampicillin, 100 μg/mL; chloramphenicol, 25 μg/mL; or kanamycin, 50 μg/mL) at 37°C with aeration. For preparation of competent cells, *E. coli* was grown in ZymoBroth (Zymo Research) at 30°C.

### Yeast strains and growth media

All *Saccharomyces cerevisiae* strains used in this study are in the S288c background, derived from the FY4 strain [83], and contain no auxotrophic or antibiotic selection markers. A complete list of strains is provided in Table S1.

Strains were constructed using markerless CRISPR/Cas9-based genome editing. Reporter constructs were integrated at the HO locus. Gene expression constructs were integrated at the AGA1 locus and the AGA2 locus. Native locus gene-tagging strains used in Figure S1A were generated using a modified C-SWAT method. Additional details of strain construction are provided in the Method Details section.

All yeast cultures were grown at 30°C. YEPD medium (1% [w/v] yeast extract, 2% [w/v] peptone, 2% [w/v] glucose) was used for preparation of cells for transformation. YEPD agar plates supplemented with antibiotics (G418, 200 μg/mL; clonNAT, 100 μg/mL; or hygromycin B, 200 μg/mL) were used for selection of transformants. For strains unable to grow on glucose (e.g., hexokinase triple mutants and hexose transporter mutants), YEPGal medium (identical to YEPD except with 2% [w/v] galactose replacing glucose) was used.

Glucose repression response measurements were performed in S(LF) medium (low-fluorescence synthetic minimal medium containing 1.7 g/L yeast nitrogen base without amino acids, riboflavin, or folic acid, and 5 g/L ammonium sulfate) supplemented with varying glucose concentrations. Cells were first inoculated from single colonies and grown to saturation in YEPD (∼24 h), followed by a second outgrowth in S(LF) medium containing 128 mM glucose for ∼16 h to an OD of ∼0.2. Cultures were then washed with S(LF) medium without glucose, and diluted into assay conditions and grown in 500 μL medium in 1.3 mL deep-well plates at 30°C with shaking at 999 rpm (Infors HT Multitron). For strains unable to grow on glucose, 128 mM galactose was used in place of glucose during outgrowth steps.

Growth curves were measured in S(LF) medium containing 128 mM glucose following an outgrowth to saturation in YEPD (∼24 h). Cells were cultured in 3 mL medium in glass culture tubes tilted at 30° from vertical and shaken at 230 rpm.

Glucose consumption rate measurements were performed in S(LF) medium containing 10 mM or 16 mM glucose following overnight growth to saturation in YEPD.

All YEP and S(LF) media used in quantitative experiments were sterilized by filtration. All S(LF) media were stored at 4°C and warmed up to room temperature before experiments.

## METHOD DETAILS

### Plasmid construction

#### Plasmids were constructed using a combination of Golden Gate assembly and Gibson assembly

Plasmid design followed the MoClo Yeast Toolkit (MoClo-YTK) standard [84]. In this study, parts spanning Type 3 and Type 4a (referred to as Type34a) were used as the standard format for coding sequences (CDSs), while Type 4b parts were used for terminators. This convention is compatible with the original MoClo-YTK system.

Additional part plasmids were generated either from synthesized gene fragments or by subcloning from existing plasmids. The NatR selection marker from the MoClo-YTK collection was avoided, as 2μ plasmids containing this version of NatR were associated with growth defects under certain conditions during yeast transformant isolation. Instead, a natMX cassette was used, subcloned from a previous plasmid (A05V).

To facilitate compatibility between Golden Gate and Gibson assembly workflows, placeholder part plasmids were introduced. For example, a Type 2 (promoter) placeholder (pPlaceholder) was assembled with an mVenus CDS to generate a pPlaceholder-mVenus construct, which served as a backbone for subsequent Gibson assembly. The placeholder region was excised using BbsI-HF (NEB), gel-purified, and replaced with a promoter sequence (e.g., pHXK1) via Gibson assembly to generate reporter constructs (e.g., pHXK1-mVenus). Similarly, Type 3 (partial CDS) placeholders (Placeholder) were used to construct backbone plasmids such as pTetO7-Placeholder-mScarlet-I for insertion of coding sequences. This strategy enabled insertion of arbitrary sequences without the need to remove internal restriction sites.

For plasmid assembly, part plasmids were prepared using spin-column miniprep (QIAprep Spin Miniprep Kit) and normalized to 40 nM. Synthetic DNA fragments (IDT gBlock or gBlocks HiFi) were resuspended in IDTE buffer (10 mM Tris, 0.1 mM EDTA; IDT) and quantified using a NanoDrop spectrophotometer. PCR products were purified by gel extraction (NEB Monarch DNA Gel Extraction Kit). During purification, care was taken to fully wet the side wall of spin column during wash steps to ensure purity.

Golden Gate reactions (10 μL total volume) contained 10 fmol backbone plasmid (0.5 μL), 20 fmol of each plasmid or PCR insert (1 μL), 1 μL T4 DNA ligase buffer (NEB), 0.5 μL T4 DNA ligase (400 U/μL, NEB M0202S), and 0.5 μL BsaI-HFv2 or BsmBI-v2 (NEB). When gBlocks were used, 30 fmol of each insert was included. To improve pipetting accuracy, DNA mixes were prepared at 2× volume and split prior to addition of the reaction master mix.

Golden Gate reactions were thermocycled as follows: for BsaI assemblies, 25 cycles of 37°C for 2 min and 16°C for 5 min, followed by 55°C for 10 min and hold at 4°C; for BsmBI assemblies, 15 cycles of 42°C for 5 min and 16°C for 5 min, followed by 55°C for 10 min and hold at 4°C. Reaction products were used immediately for transformation into *E. coli* or stored at −20°C.

For insertion of annealed oligonucleotides (e.g., sgRNA constructs), 1 μL each of 100 μM forward and reverse oligos were mixed and diluted to 20 μL in water. Subsequently, 5 μL of this mixture was diluted to 50 μL in water or STE buffer (10 mM Tris pH 8.0, 50 mM NaCl, 1 mM EDTA). Oligos were annealed by heating to 96°C for 6 min followed by cooling to 23°C at 0.1°C/s. Annealed oligos (0.5 μL) were used per 10 μL Golden Gate reaction.

For ligation of multiple oligo pairs (e.g., synthetic promoter construction), each oligo was first phosphorylated in a 10 μL reaction containing 1 μL 100 μM oligo, 1 μL T4 DNA ligase buffer (NEB), and 1 μL T4 polynucleotide kinase (NEB). Complementary oligos were then combined, diluted, and annealed as described above.

Gibson assembly was performed following Gibson et al. [85]. Reactions (15 μL) contained 25 fmol backbone DNA and 50 fmol of each insert and were incubated at 50°C for 60 min. A 1 μL aliquot of the reaction was used for transformation into *E. coli*.

### Bacterial transformation

Chemically competent *E. coli* cells were prepared using the Mix & Go! *E. coli* Transformation Kit (Zymo Research, T3001). A single colony was grown overnight to saturation in 5 mL LB medium at 37°C. The overnight culture was diluted 1:100 into 100 mL ZymoBroth in a 500 mL flask and grown at 30°C. After ∼6 h, when the culture reached an OD_600_ of 0.4–0.6 (target ∼0.55), cells were chilled on ice for 10 min.

Wash and competent buffers were prepared according to the manufacturer’s instructions. All buffers, tubes, and pipetting materials were pre-cooled to 4°C.

Cells were harvested by dividing the culture into four 50 mL tubes and centrifuging at 1,600 × g for 10 min at 4°C. After removal of the supernatant, each pellet was resuspended in 2.5 mL wash buffer. Suspensions were combined into two tubes and incubated on ice for 10 min. Cells were then pelleted again under the same conditions, and each pellet was resuspended in 5 mL competent buffer. Suspensions were combined into a single tube and incubated on ice for an additional 10 min.

Aliquots of 115 μL or 230 μL were dispensed into 1.5 mL microcentrifuge tubes using a repeater pipette (Eppendorf Repeater E3) at low speed. Cells were flash-frozen in liquid nitrogen or a dry ice–ethanol bath and stored at −80°C.

For transformation, competent cells were thawed on ice for 15–30 min and gently resuspended by flicking. For each transformation, 25 μL of competent cells was mixed with 0.5–2 μL DNA and incubated on ice for 30 min. Cells were heat shocked at 42°C for exactly 30 s using a pre-heated heat block (with water added to wells to improve thermal transfer), then returned to ice for 1 min. SOC medium (room temperature) was added, and cells were recovered at 37°C for 1 h with shaking. Transformation mixtures were plated onto pre-warmed LB plates with the appropriate antibiotic using glass beads and incubated at 37°C overnight.

### Yeast transformation

Yeast transformation was performed using a standard lithium acetate/single-stranded carrier DNA/PEG protocol [86, 87].

For preparation of fresh competent cells, yeast colonies were grown overnight to saturation in YEPD medium at 30°C, diluted to OD_600_ = 0.1, and grown for ∼4 h to OD_600_ ≈ 0.4. For slow-growing strains, a higher starting OD (e.g., 0.15) was used. Cells corresponding to 2.0 mL at OD_600_ = 0.4 per transformation were collected by centrifugation at 3,000 × g for 3 min, washed once with 10 mL sterile water, and resuspended in 100 mM lithium acetate to a final volume of 100 μL per transformation.

For experiments requiring repeated transformations, frozen competent yeast cells were prepared. Overnight cultures were diluted to OD_600_ = 0.1 in 50 mL YEPD and grown to OD_600_ = 0.7–1.0 (target ∼0.9). Cells (40 mL) were harvested by centrifugation at 3,000×*g* for 3 min at room temperature, washed with 10 mL sterile water, and pelleted again. The pellet was resuspended in 200 μL sterile water (0.005× culture volume), mixed with an equal volume (200 μL) of 20% glycerol. Aliquots (85 μL) were dispensed into microcentrifuge tubes and frozen gradually (by placing in a styrofoam container) at −80°C.

For use, frozen competent cells were thawed at room temperature for ∼5 min, pelleted at 16,000 × g for 30 s, and resuspended in 100 μL 100 mM lithium acetate per 25 μL frozen aliquot.

For transformation, 100 μL of competent cells (fresh or thawed) was pelleted at 16,000×*g* for 30 s and resuspended in 20 μL DNA mixture. A total of 124 μL transformation master mix was added (per reaction: 96 μL 50% PEG 3350, 14.4 μL 1 M lithium acetate, 4 μL boiled ssDNA, and 9.6 μL water). ssDNA (55 μL per aliquot) was boiled at 99°C for 5 min and immediately cooled on ice prior to use. The mixture was vortexed thoroughly, incubated at 30°C for 30 min, vortexed again, and heat shocked at 42°C for 15 min.

Following heat shock, cells were pelleted at 8,000×*g* for 1.5 min, resuspended in 800 μL YEPD, and recovered in culture tubes at 30°C with shaking for 3 h. An appropriate volume of cells (e.g. 60 μL or 90 μL) were plated onto pre-warmed YEPD agar plates with the appropriate antibiotic using glass beads and incubated at 30°C for ∼2 days. Comparable numbers of transformants were observed for fresh and frozen competent cells protocols.

Because individual colonies may contain mixed populations at a non-negligible rate, three colonies from each transformation plate were restreaked onto non-selective YEPD agar plates and grown for 2 days for single colonies. Colonies were genotyped by colony PCR or sequencing where appropriate and subsequently cultured in YEPD for phenotypic analysis and glycerol stock preparation. Typically, over 90% of colonies passed validation.

### Markerless genome editing with CRISPR/Cas9

#### Genome editing was performed using markerless CRISPR/Cas9-assisted homologous recombination

Cas9 and sgRNA were delivered using a two-plasmid system [88]. Cas9 was expressed from the pRPL18B promoter on a 2μ plasmid carrying either KanR, NatR, or HygR selection markers. sgRNAs were expressed as yeast tRNA–HDV ribozyme fusions under control of yeast tRNA promoters. Prior to transformation, Cas9 plasmids and sgRNA plasmids were linearized using BsaI-HFv2 and EcoRV-HF (NEB), respectively. Linearized Cas9 plasmids were gel-purified (NEB Monarch DNA Gel Extraction Kit), whereas sgRNA plasmids were purified using a spin-column cleanup kit (Qiagen QIAquick PCR Purification Kit). The resulting fragments contained homologous regions enabling in vivo assembly by homologous recombination into a single plasmid carrying both Cas9 and sgRNA expression cassettes. This strategy enabled flexible pairing of sgRNAs with different selectable markers without additional cloning steps.

sgRNAs were designed to minimize predicted off-target activity and to achieve Azimuth-predicted [89] on-target scores above 0.7. sgRNA sequences were synthesized as complementary oligonucleotides and cloned into the sgRNA entry vector pWS082.

For each transformation, donor DNA (120 fmol linear DNA fragments) was co-transformed with linearized Cas9 plasmid fragments (40 ng) and sgRNA plasmid fragments (80 ng). Transformants were selected using the corresponding antibiotics (G418 for KanR, clonNAT for NatR, or hygromycin B for HygR). Following strain isolation, Cas9 plasmids were readily lost during growth in non-selective YEPD medium, resulting in markerless edited strains.

For repeated editing at commonly used genomic loci, a one-plasmid Cas9/sgRNA system was used to simplify plasmid preparation. We observed that transformation efficiency of linear DNA fragments was substantially higher (∼10-fold) than that of supercoiled plasmids. Therefore, plasmids were designed to be linearized prior to transformation and re-circularized in vivo by homologous recombination. To enable this, selectable marker coding sequences were split into overlapping fragments separated by an EcoRV site. For example, the split KanR cassette (referred to as KanRS) consisted of KanR(1–501 bp), EcoRV site and KanR(301–810 bp). Plasmids were linearized with EcoRV, purified using a spin-column cleanup kit (Qiagen QIAquick PCR Purification Kit), and 40 ng DNA was used per transformation.

For genomic integrations, donor plasmids containing homology regions were linearized with restriction enzymes (typically NotI-HF, depending on construct design), purified using a spin-column cleanup kit (Qiagen QIAquick PCR Purification Kit), and used as homologous recombination templates. Three genomic integration sites were used in this study: the HO locus (replacing −1814 bp to +1198 bp), the AGA1 locus (replacing −366 bp to +412 bp), and the AGA2 locus (replacing −424 bp to +87 bp). Coordinates are reported relative to the ATG start codon, where the A is defined as +1 and the preceding nucleotide as −1.

For gene deletions, donor DNAs were generated by annealing and extending two oligonucleotides containing a 20 bp complementary region and 40–60 bp homology arms flanking the target locus. Extension reactions were performed using Q5 polymerase (NEB) without template DNA for three PCR cycles, and 2 μL reaction product was used per transformation. The central 20 bp region contained a unique deletion barcode including a PAM sequence, allowing the deleted locus to be retargeted by CRISPR/Cas9 for subsequent genome editing. Gene replacement with mutant alleles was performed by first generating a deletion strain and subsequently targeting the barcode sequence for reintegration of the desired mutant.

For N- or C-terminal fluorescent protein tagging, sgRNAs were designed such that insertion of the fluorescent protein coding sequence disrupted the sgRNA target site upon successful integration by splitting the target sequence into two nonfunctional halves.

### Gene tagging with modified C-SWAT

When suitable sgRNA target sites were not available near the C-terminus of a gene, gene tagging was performed using a modified C-SWAT approach [90].

A modified C-SWAT module was constructed consisting of a GDGAGS linker, the N-terminal fragment of mScarlet-I (bp 1–240), an ADH1 terminator, a hygromycin resistance marker (HygR), an I-SceI recognition site, and the C-terminal fragment of mScarlet-I (bp 457–699) (linker-mScarlet-I(1-240)-tADH1(180 bp)-HygR-I-SceI-mScarlet-I(457-699)). This design avoids internal sequence overlap, allowing specific amplification under standard PCR conditions.

For genomic integration, the C-SWAT module was amplified using Q5 polymerase (NEB) with primers introducing homology arms (40 bp or 60 bp) targeting insertion immediately upstream of the endogenous terminator via homologous recombination. Yeast transformation was performed with the PCR product, and transformants were genotyped by colony PCR to confirm correct integration. Sequencing was performed when needed.

To remove the selection marker and restore a full-length mScarlet-I coding sequence, a second transformation was performed. Cells were transformed with a plasmid expressing I-SceI (CEN/ARS backbone, I-SceI under the pTEF2 promoter) together with a homologous recombination repair template corresponding to the internal region of mScarlet-I (from 41 to 656 bp). Two independent isolates from the initial integration were used for this step. Following transformation, colonies were screened for fluorescence, yielding marker-free, fluorescently tagged strains. The CEN/ARS plasmid was subsequently lost during growth in non-selective YEPD medium.

This approach enabled markerless C-terminal tagging of genes while preserving native terminators.

### Glucose response measurement

To measure gene expression or localization responses across different glucose concentrations, cells were cultured in S(LF) medium supplemented with varying glucose concentrations following sequential outgrowth steps in YEPD and S(LF) + 128 mM glucose medium.

For the initial YEPD outgrowth, single colonies were picked using 20 μL pipette tips and resuspended in 50 μL water in a PCR plate. Aliquots of 5 μL cell suspension (or up to 20 μL for slow-growing strains) were used to inoculate 500 μL YEPD medium in 1.3 mL 96-well deep-well plates. Cultures were grown at 30°C with shaking at 999 rpm in a humidified incubator (80% relative humidity; Infors HT Multitron).

After ∼24 h growth, saturated YEPD cultures were used to inoculate an S(LF) + 128 mM glucose outgrowth plate. For each strain, 5 μL saturated culture was diluted 1:160 into 800 μL S(LF) + 128 mM glucose medium. Cultures were then serially diluted 2-fold to generate final dilutions of 1:320, 1:640, 1:1280, 1:2560, 1:5120, and 1:10240, each in 400 μL medium. Plates were incubated under the same shaking conditions.

Following ∼16–17 h growth, OD_600_ values of the outgrowth cultures were measured to identify wells in the appropriate growth range. For measurement, 150 μL culture was transferred to clear flat-bottom 96-well plates (CellTreat 229197), and OD_600_ was measured using a BioTek Synergy plate reader. Measurements were converted to equivalent 10 mm path length OD_600_ values using a calibration curve. For each strain, the dilution with the highest OD_600_ below 0.27 was selected for subsequent glucose response measurements. Selected wells typically exhibited OD_600_ values between 0.1 and 0.27 and corresponded to 1:2560, 1:5120, or 1:10240 dilutions.

The volume of each selected culture required to achieve OD_600_ = 0.2 in a final volume of 100 μL was calculated. The corresponding volume of culture was collected, washed twice with 400 μL glucose-free S(LF) medium by centrifugation (3,000 × g, 3 min) followed by resuspension, and finally resuspended in 100 μL glucose-free S(LF) medium. Improved recovery during wash steps was observed when using repeatedly autoclaved deep-well plates or low protein-binding plates; repeatedly autoclaved deep-well plates were used throughout this study.

For glucose titration experiments, a 1.25× glucose titration plate was prepared in 1.3 mL deep-well plates containing 400 μL S(LF) medium supplemented with 1.25× final glucose concentrations (e.g., 160 mM, 80 mM, …, 1.25 mM). To inoculate the titration plate, 50 μL normalized culture was first diluted 1:50 into 1,000 μL glucose-free S(LF) medium in a 2 mL deep-well plate. Aliquots of 100 μL diluted culture were then dispensed into each titration well, yielding final cultures of 500 μL with 0.002 OD containing glucose concentrations ranging from 128 mM to 1 mM.

Following inoculation, cultures were incubated under the same growth conditions for ∼7 h prior to microscopy measurements.

### Multiplexed glucose response measurement

For multiplexed glucose repression measurements using mixed strain populations, OD_600_ measurement, normalization, and mixing steps were automated using a Tecan Fluent 1080 liquid handler to reduce experimental time and minimize manual error.

OD_600_ measurements of outgrowth cultures were performed by transferring 150 μL culture into clear flat-bottom plates using a 96-channel pipetting head (Tecan MCA 96). OD_600_ measurement and selection of appropriate outgrowth wells were performed as described for non-multiplexed glucose response measurements.

After selecting wells within the desired OD range, cultures were resuspended and normalized during the mixing process using a multi-aspiration single-dispense workflow controlled through a custom Python script interfacing with the Tecan API via ExecutionChannel. To generate a mixed population, a single 1,000 μL tip sequentially aspirated appropriate volumes from each of nine cultures and dispensed the combined mixture into the destination well using the Tecan Flexible Channel Arm (FCA). Mixed cultures were washed and normalized to OD_600_ = 0.4 in 200 μL glucose-free S(LF) medium. Subsequently, 50 μL of the mixed culture was used for glucose titration inoculation following the same protocol described above for non-multiplexed experiments.

The Python API code used for Tecan Fluent control is available at https://github.com/angli339/tecan-fluent-api.

### High-throughput fluorescent microscopy

Concanavalin A (ConA)-coated glass-bottom 384-well plates were used for microscopy. ConA solution was prepared as a 5× stock following Hansen et al. [91] and stored at −80°C. The 1× working solution contained 4 mg/mL ConA, 10 mM CaCl₂, and 10 mM MnCl₂ in 0.22× PBS (pH 6.5). After dilution to 1×, the solution was stored at 4°C and remained stable for at least one month. Precipitates may form during storage at 4°C and were removed by filtration without affecting performance.

For coating, 50 μL ConA was dispensed into alternating wells of the first row and sequentially transferred across wells using a multichannel pipette. Care was taken to fully wet the glass surface during each transfer. Each 50 μL aliquot was used to coat 16 wells. Immediately after coating, wells were then washed once with Milli-Q water by adding 70 μL per well and aspirating using an 8-channel manifold (BrandTech 704526) fitted with 10 μL tips. Plates were incubated at 30°C for ∼30 min to allow drying and pre-warming. To limit imaging time, experiments typically used no more than half of a 384-well plate.

Before imaging, 90 μL of cell culture from deep-well plates was transferred into the ConA-coated 384-well plate. Plates were centrifuged at 50×g for 2 min to allow cells to settle to the plate bottom. To protect the glass surface from contamination from dust, a cleanroom wipe was placed beneath the plate during centrifugation.

For imaging with an oil-immersion objective, immersion oil (Olympus IMMOIL-F30CC) was applied to the underside of the plate by holding the plate horizontally and dispensing oil directly onto the glass surface from below.

Imaging was performed using a motorized Nikon Ti-E inverted microscope equipped with a motorized XY stage (Prior ProScan II), a metal halide light source with excitation filter wheel and tunable shutter (Prior Lumen 200), an emission filter wheel (Prior Scientific HF110), and a CCD camera (Hamamatsu ORCA-R2). A 60× oil-immersion objective (Plan Apo VC 60×/1.40 NA) was used. Excitation and emission filters (Semrock) for each channel were as follows: BFP (Ex 390/40, Em 452/45), CFP (Ex 438/24, Em 483/32), YFP (Ex 500/24, Em 542/27), and RFP (Ex 562/40, Em 641/75). A multi-band dichroic beamsplitter (Semrock FF444/520/590-Di01) was used for all channels. Illumination intensity was optimized to minimize photobleaching, with typical exposure times ranging from 100 to 500 ms. A 2×2 binning mode was used for FRET sensor and Mig1 localization experiments.

Image acquisition, visualization, and quantification were performed using custom microscope control software written in C++/Qt. A Python script (Jupyter Notebook) was used for experimental setup and high-level control of acquisition logic. Source code of the control software is available at https://github.com/angli339/NikonTiCtrl.

During acquisition, bright-field images were used for real-time cell segmentation using a custom U-Net model. Cell counts were accumulated for each well, and the number of imaging sites was dynamically adjusted based on cell density. Imaging positions were arranged in a non-overlapping grid. For standard experiments, at least 400 cells per well were targeted in no less than 6 imaging sites. For multiplexed experiments, a minimum of 1,800 cells per well was targeted.

### Training of finetuned U-Net model

A 2D U-Net architecture based on Falk et al. [92] was used as the base model for fine-tuning. The model was reimplemented in TensorFlow with the modification that zero-padding was applied to Conv2D layers (`padding=’same’`), ensuring that the output segmentation maps have the same spatial dimensions as the input images. Pretrained weights (2d_cell_net_v0) were obtained from https://lmb.informatik.uni-freiburg.de/ resources/opensource/unet/ and converted from Caffe format to TensorFlow using a custom conversion function.

To generate accurate ground truth annotations, genetically encoded fluorescent markers were used to label cellular structures. Whole cells were labeled with mCherry expressed under the constitutive pTDH3 promoter. Nuclei were labeled with mTagBFP2 fused to an N-terminal BPSV40 nuclear localization signal [93] and expressed under the pACT1 promoter. The plasma membrane was labeled using mNeonGreen fused to the C-terminal palmitoylation domain of Gap1 (residues 572–602) [94], expressed under the pACT1 promoter. This Gap1-derived sequence reliably targeted fluorescent proteins to the plasma membrane in yeast.

Bright-field, BFP, YFP, and RFP images were acquired at both low cell density (sparse distribution) and high cell density (crowded fields with out-of-focus cells). Initial segmentation masks were generated using classical image processing methods. Specifically, a Laplacian-of-Gaussian filter was applied to the plasma membrane channel for edge detection, followed by thresholding, removal of small objects, and morphological closing. These masks were manually curated in Napari to remove out-of-focus cells and separate contacting cells. Bright-field, nuclear, and cytoplasmic channels were used as references during manual curation to determine cell focus and boundaries of contacting cells.

A total of nine annotated images (seven at typical cell density and two at high density) were used as ground truth for training. Three additional images were used for validation. Model performance was primarily evaluated by visual inspection to minimize the need for extensive manual annotation. To improve robustness to non-cellular artifacts, an additional nine training images from independent experiments were included by manually correcting segmentation outputs from a previous model version and removing non-cell objects.

Segmentation masks were defined such that pixel values of 1 correspond to cell interiors, whereas 0 correspond to background and cell–cell boundaries. This labeling scheme ensures that each connected region represents a single cell and trains the model to explicitly identify boundaries between contacting cells.

Training was performed on a single NVIDIA GTX 1070 GPU (8 GB memory). Bright-field images were normalized to the range [0, 1] using min–max scaling. Data augmentation included random zoom (±20%), random flipping, and random cropping to 512 × 512 pixels. To improve performance on binned data, simulated 2×2 binned images and corresponding segmentation masks were included during augmentation. Models were trained in float32 precision using the Adam optimizer (learning rate = 1 *×* 10*^−^*^5^) and binary cross-entropy loss. Noticeable improvement in segmentation quality was observed after one epoch, and training was continued for a total of nine epochs. The trained model and weights are available at https://github.com/angli339/unet_yeast_bf and https://huggingface.co/angli339/unet_yeast_bf.

Inference was performed in float16 precision without noticeable loss in accuracy. The model was deployed using TensorFlow Serving for real-time segmentation during image acquisition.

Benchmarking of models was performed on a system equipped with an Intel Xeon E3-1245 v5 CPU and an NVIDIA GTX 1070 GPU, using input images of size 1344 × 1024 pixels. Segmentation speed and accuracy were compared with Cellpose [95] and YeaZ [96], with results reported in Figure S3.

### Image segmentation and quantification

Bright-field images were normalized to the range [0, 1] using min–max scaling and used as input to a U-Net model for segmentation. For low-resolution images acquired with 2×2 binning, images were upscaled by a factor of 2 prior to segmentation, and the resulting segmentation output was subsequently downscaled to the original resolution. Pixel-wise segmentation scores were thresholded at 0.5 to generate an initial binary mask. For each connected region, the mean segmentation score was calculated, and regions with an average score below 0.95 were discarded to remove low-confidence non-cell objects.

Because the U-Net model was trained to separate contacting cells, no additional post-processing was required for cell separation. Segmentation results were manually inspected for at least two representative images per experiment to ensure quality. This pipeline effectively excludes out-of-focus cells and is robust to morphological variation, including elongated cell shapes.

Cellular fluorescence was quantified by calculating the mean pixel intensity within each segmented region for each fluorescence channel. After subtracting the camera’s offset (baseline signal in the absence of light), mean pixel intensities were normalized by exposure time and converted to physical units using the camera conversion factor (0.28 electrons per AD count, calculated for 16-bit readout based on the camera’s full-well capacity and offset). The resulting values are reported in units of electrons per pixel per millisecond. For images acquired with 2×2 binning, values were divided by 4 to obtain the pre-binning photoelectron flux per pixel.

Illumination intensity was not normalized across experiments and may vary over time or between acquisition settings. Therefore, direct comparison of fluorescence intensities between experiments was avoided unless normalization could be achieved using biological controls.

Following single-cell quantification, fluorescence intensity distributions were visualized as histograms for each channel to assess population-level behavior and data quality. Histograms were evaluated manually for quality control. Population-average fluorescence values were calculated in log space and are reported in the figures throughout this study.

### Growth curve measurement

Following ∼24 h of outgrowth in YEPD medium containing 2% glucose, cultures were diluted 1:2000 into 3 mL S(LF) medium supplemented with 128 mM glucose (2.3%) in 16 mm diameter glass culture tubes (Corning 9820-16). Cultures were incubated at 30°C with shaking at 230 rpm. Optical density at 600 nm (OD_600_) was recorded every 30 s using a custom growth measurement device equipped with an array of eight LEDs and eight photodiodes configured for transmission measurements. A detailed description of the device will be provided in a forthcoming manuscript. Measurements were continued until cultures reached saturation. Raw OD_600_ values were converted to linearized equivalent 10 mm path length OD_600_ using a standard curve.

To ensure accurate blanking, culture tubes were prefilled with 2.9 mL medium using a repeater pipette (Eppendorf Repeater E3) and placed into the measurement device for baseline recording. Without removing the tubes, 0.1 mL of diluted inoculum was added to each tube using a multichannel pipette, and data acquisition was continued.

For reuse, glass culture tubes were decontaminated with 10% bleach and cleaned by overnight soaking in 5% Contrad 70 (Decon Labs). Tubes were then thoroughly rinsed with deionized water and dried upside down on cleanroom wipes. A thin layer of mineral oil was applied to the outer surface near the opening to facilitate reattachment of tube closures (Bal Supply 16144CL). Tubes were sterilized by autoclaving at 121°C for 20 min. Stainless steel racks (Utile Scientific 4050.4605.2) were used during autoclaving to prevent residue formation associated with certain plastic racks.

Growth curves were processed by applying a median filter (kernel size = 5) to remove occasional spikes in raw data. Doubling times were calculated using a 60 min sliding window to assess stability during exponential growth. Reported doubling times correspond to the average value measured over the OD_600_ range of 0.05–0.2. All measurements were performed in biological triplicate, and individual replicates are shown in figures to illustrate reproducibility.

### Glucose consumption rate measurement

Saturated YEPD cultures were diluted 1:6400 into 15 mL S(LF) medium containing 16 mM glucose. For doxycycline titration experiments, media were additionally supplemented with doxycycline at concentrations ranging from 0.02 to 1.28 μg/mL. Cultures were grown for 16 h at 30°C with shaking, after which OD_600_ was measured using a spectrophotometer. Cultures were then diluted in the same medium to OD_600_ = 0.05 (15 mL total volume) and glucose consumption rate measurement was started.

Samples were collected every 30 min over a 6 h time course. At each time point, 0.6 mL was removed for OD_600_ measurement, and 100 μL was transferred to a PCR plate and kept on ice.

At the end of the time course, glucose concentrations in the media were measured. Samples were equilibrated to room temperature for pipetting accuracy and 50 μL of each sample was diluted 1:20 with water to a final volume of 1,000 μL in a 2 mL deep-well plate, followed by centrifugation at 3,000 × g for 3 min. Supernatants (50 μL, in triplicate) were used for glucose quantification using the D-Glucose HK Assay Kit (Megazyme), following the manufacturer’s instructions. Reactions were performed in UV-transparent microplates (Greiner Bio-One 655809) with a total volume of 150 μL per reaction. After incubation at room temperature for 5 min, absorbance at 340 nm (OD_340_) was measured using a plate reader. A two-point standard curve (0 mM and 16 mM glucose) was used to calculate glucose concentrations.

To calculate glucose consumption rates, first, doubling time (*t*_doubling_) was fitted from OD_600_ measurements. Then, glucose concentration was plotted as a function of OD_600_, and a linear fit was used to obtain the slope (*k*, in units of mM/OD). The glucose consumption rate (mM/min/OD) was calculated as *−k ×* ln(2)/*t*_doubling_, and reported accordingly.

## QUANTIFICATION AND STATISTICAL ANALYSIS

Image analysis and fluorescence quantification are described in the Method Details section. Statistical analyses were performed using the Python scipy.stats module.

The exact value of *n*, the definition of *n*, the number and type of replicates, type of summary statistics, and statistical tests are provided in the corresponding figure legends. For most experiments, conclusions were supported by large effect sizes that were readily apparent from the measurements. Consequently, formal statistical testing was limited to straightforward comparisons of glucose response phenotypes, with results presented in Figure S11 and Figure S12. No statistical methods were used to predetermine sample size.

To minimize bias during image acquisition and analysis, field of views in microscopy were acquired automatically on a predefined non-overlapping grid without manual selection. Cell segmentation and rejection of out-of-focus cells were performed automatically using bright-field images rather than fluorescence images, preventing fluorescence intensity from influencing segmentation or cell selection for fluorescence quantification.

