## Supplementary material for "Glucose repression of *HXK1* is glucose flux-dependent via non-canonical regulation of Mig1": Document S1

Ang Li [1] and Michael Springer [1,2,3,*]

[1]Department of Systems Biology, Harvard Medical School, Boston, MA, USA
[2]Broad Institute of Massachusetts Institute of Technology and Harvard, Cambridge, MA, USA
[3]Wyss Institute for Biologically Inspired Engineering, Harvard University, Boston, MA, USA

### Supplemental Figures

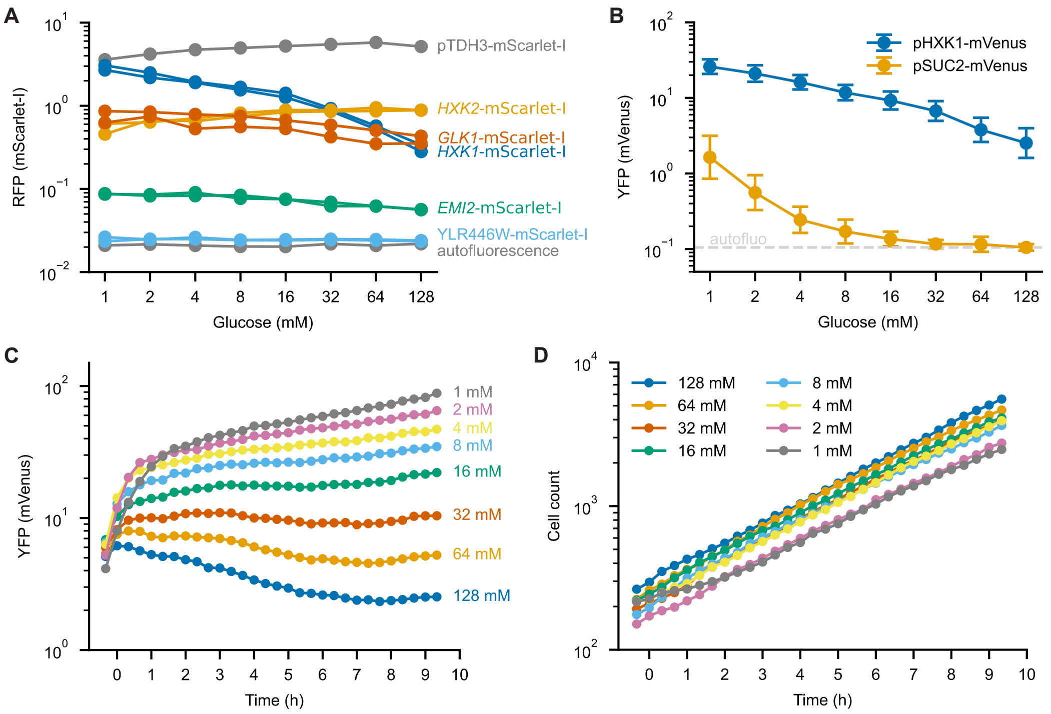


**Figure S1. Development and validation of the pHXK1 transcriptional reporter, related to Figure 1.**
**(A)** Steady-state expression of endogenous hexokinases (*HXK1* and *HXK2*), glucokinase (*GLK1*), and hexokinase-like proteins (*EMI2* and YLR446W) fused to mScarlet-I at their native loci across a glucose titration. A constitutively expressed pTDH3-mScarlet-I reporter and autofluorescence control are included for comparison. Each line represents one biological replicate ($n = 2$ for mScarlet-I fusion strains; $n = 1$ for the pTDH3-mScarlet-I and autofluorescence control strains).
**(B)** Comparison of steady-state glucose responses of the pHXK1-mVenus and pSUC2-mVenus transcriptional reporters. Points show the population mean, and error bars indicate the standard deviation of single-cell fluorescence ($n \geq 456$ cells per condition). One representative biological replicate is shown.
**(C)** Time course of pHXK1-mVenus reporter fluorescence following transfer to the indicated glucose concentrations. Cells were cultured on the microscope without shaking. Each line represents the population mean fluorescence.
**(D)** Cell counts measured from the same microscopy dataset shown in **(C)**. Cell number increased throughout the experiment at all glucose concentrations, confirming continued cell growth during the time course.

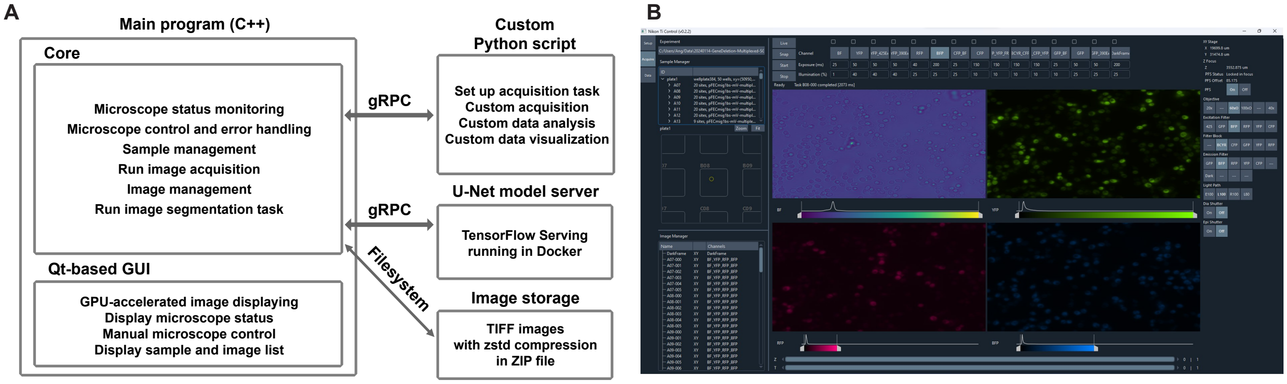


**Figure S2. Custom software for automated high-throughput fluorescence microscopy, related to Figure 1.**
**(A)** Software architecture of the custom microscope control system. The core acquisition program, written in C++ with a Qt-based graphical user interface, controls microscope operation, image acquisition, sample management, image processing, and image segmentation. Custom Python scripts communicate with the acquisition program through gRPC to define acquisition workflows and experimental logic, and can additionally be used for custom data analysis and visualization. U-Net image segmentation is performed by TensorFlow Serving running in a Docker container. Images are stored as TIFF files with Zstandard (zstd) compression in ZIP archives. The software is available open source at GitHub at https://github.com/angli339/NikonTiCtrl.
**(B)** Screenshot of the main microscope control program during image acquisition. The program is controlled by custom Python scripts, which define the acquisition tasks, experimental logic, data analysis, and visualization. The graphical interface displays live multi-channel images, microscope status, acquisition settings, sample layout, and image management, and supports manual microscope control and real-time visualization of acquired images.

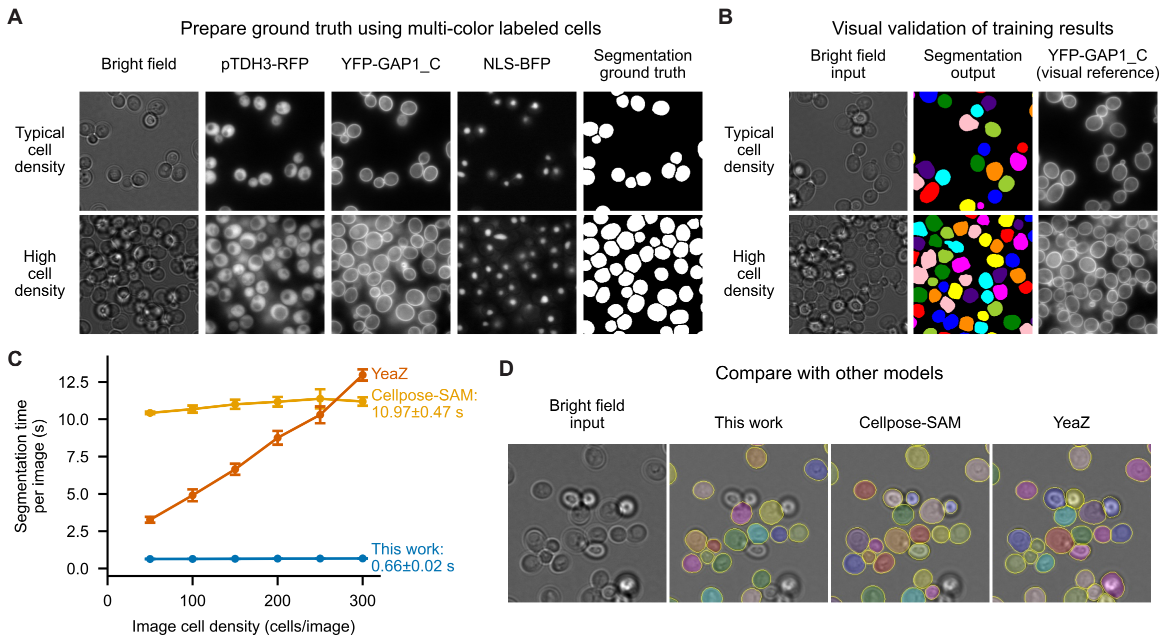


**Figure S3. Training, validation, and benchmarking of the U-Net image segmentation model, related to Figure 1.**

**(A)** Preparation of ground-truth segmentation masks for model training. Yeast cells expressing cytoplasmic mCherry, plasma membrane-localized mNeonGreen fused to the Gap1 C-terminal palmitoylation domain (Gap1 residues 572–602), and nuclear-localized mTagBFP2 carrying an N-terminal BP-SV40 nuclear localization signal were imaged together with bright-field images. Initial segmentation masks were generated from the plasma membrane channel using classical image-processing methods and manually curated using all fluorescence channels to remove out-of-focus cells and separate contacting cells. Representative images at typical and high cell densities are shown.

**(B)** Representative segmentation results of the trained U-Net model. Bright-field images were used as model input, and the plasma membrane channel is shown only as a visual reference for evaluating segmentation accuracy. Representative results are shown for images with typical and high cell densities.

**(C)** Segmentation speed of the U-Net model ("This work") compared with Cellpose-SAM and YeaZ as a function of image cell density. Benchmarking was performed using 1344 × 1024 pixel bright-field images on a system equipped with an Intel Xeon E3-1245 v5 CPU and an NVIDIA GTX 1070 GPU. Points show the mean segmentation time, and error bars indicate the standard deviation across benchmark images.

**(D)** Representative segmentation results of the U-Net model ("This work") compared with Cellpose-SAM and YeaZ on the same bright-field image. Colored regions indicate individual segmented cells. The U-Net model developed in this work selectively segments in-focus cells while rejecting out-of-focus cells, which is advantageous for quantitative measurements of cellular fluorescence.

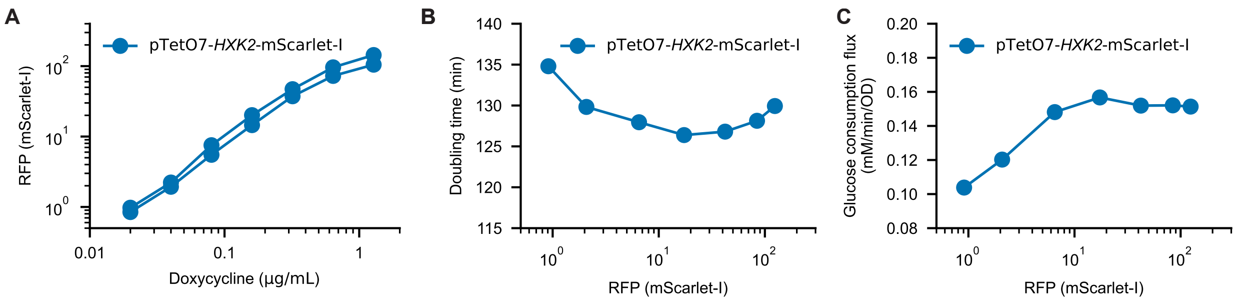


**Figure S4. Titration of HXK2 expression enables modulation of glucose consumption flux, related to Figure 2.**
*HXK1* and *HXK2* were deleted, and *HXK2*-mScarlet-I was expressed from the doxycycline-inducible pTetO7 promoter. Cells were grown in media containing 16 mM glucose.

**(A)** Hxk2-mScarlet-I expression as a function of doxycycline concentration. Each line represents one biological replicate (*n* = 2).

**(B)** Doubling time determined from $OD_{600}$ growth measurements during the glucose depletion assay as a function of Hxk2-mScarlet-I expression level. One biological replicate is shown.

**(C)** Glucose consumption flux as a function of Hxk2-mScarlet-I expression level. Glucose consumption flux was calculated from the glucose depletion rate and doubling time. One biological replicate is shown.

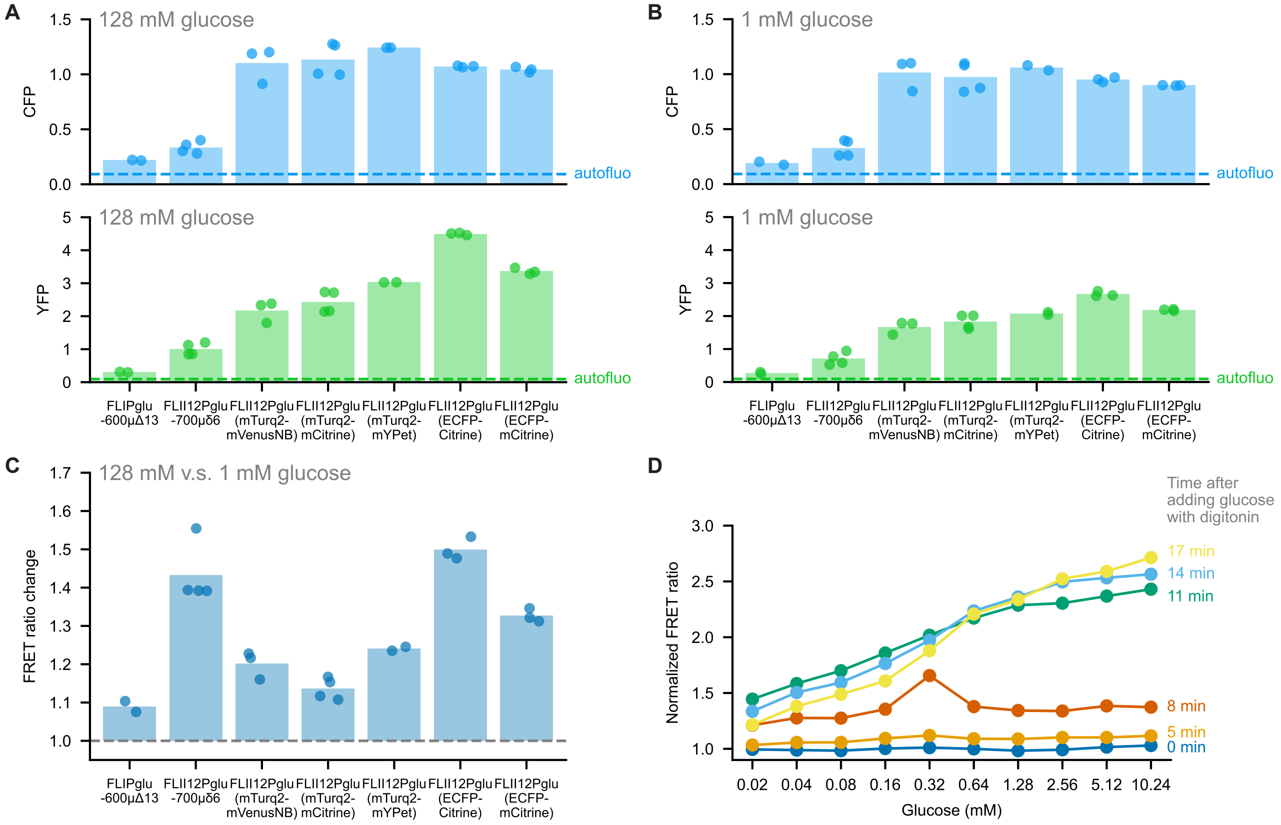


**Figure S5. Optimization and validation of glucose FRET sensor FLII$^{12}$Pglu, related to Figure 2.**
**(A–C)** *In vivo* characterization of FLII$^{12}$Pglu glucose FRET sensor variants at steady state. Cells expressing the original FLIPglu-600μΔ13 and FLII$^{12}$Pglu-700μδ6 sensors, together with FLII$^{12}$Pglu variants, were cultured in either 128 mM **(A)** or 1 mM **(B)** glucose. Mean CFP (top) and YFP (bottom) fluorescence intensities are shown in **(A)** and **(B)**; dashed lines indicate the autofluorescence background. **(C)** shows the corresponding FRET response, calculated as the ratio of the YFP/CFP fluorescence ratios in cells grown in 128 mM versus 1 mM glucose. Dots represent individual biological replicates ($n \geq 2$), and bars indicate the mean.
**(D)** Response of the optimized FLII$^{12}$Pglu(ECFP-Citrine) sensor to glucose in digitonin-permeabilized cells. Cells were grown in 128 mM glucose, washed with glucose-free medium, and incubated with the indicated glucose concentrations in the presence of digitonin to permeabilize the plasma membrane. The FRET ratio was measured over time and normalized to the FRET ratio at 0 mM glucose. The sensor response reached a near-steady state approximately 11 min after permeabilization.

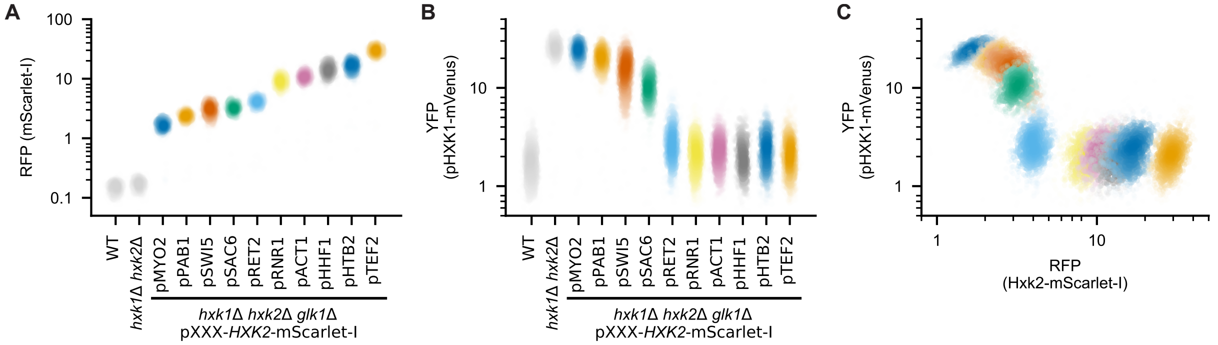


**Figure S6. Titration of *HXK2* expression using constitutive promoters in an *hxk1*Δ *hxk2*Δ *glk1*Δ strain, related to Figure 2.**

**(A)** Hxk2-mScarlet-I expression driven by a series of constitutive promoters.

**(B)** pHXK1-mVenus fluorescence measured in the same strains shown in **(A)**.

**(C)** Relationship between Hxk2-mScarlet-I abundance and pHXK1-mVenus fluorescence obtained by combining the single-cell measurements shown in **(A)** and **(B)**.

Each point represents one cell; $n \geq 530$ cells per strain. Two biological replicates were performed, data from one representative replicate are shown.

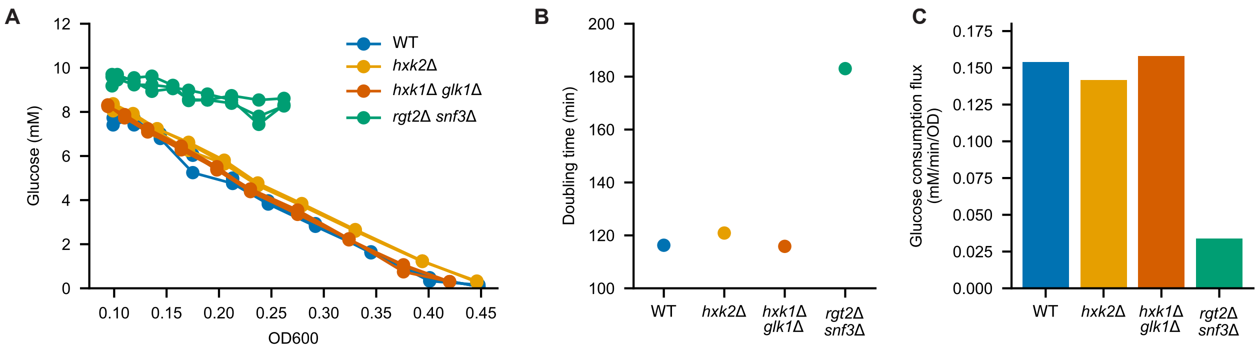


**Figure S7. Glucose consumption flux of the *snf3*Δ *rgt2*Δ mutant, related to Figure 3.**
**(A)** Glucose depletion during the glucose consumption flux assay. Glucose concentration is plotted as a function of cell density ($OD_{600}$). Each line represents one technical replicate ($n = 3$).
**(B)** Doubling times determined from $OD_{600}$ growth measurements during the glucose depletion assay in **(A)**. One biological replicate is shown.
**(C)** Glucose consumption flux calculated from the glucose depletion measurements in **(A)** and the doubling times in **(B)**. One biological replicate is shown.
Cells were grown in media containing 10 mM glucose.

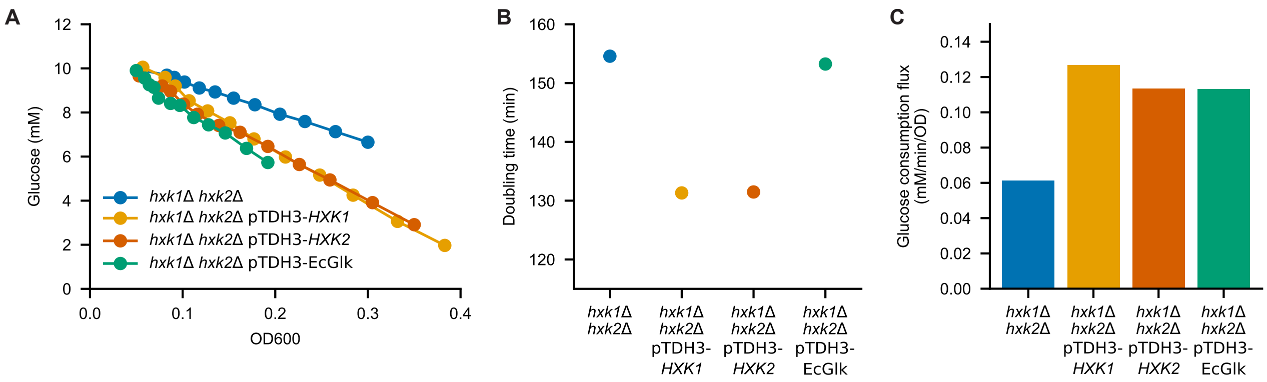


**Figure S8. Glucose consumption flux of hexokinase replacement strains, related to Figure 4.**

**(A)** Glucose depletion during the glucose consumption flux assay. Glucose concentration is plotted as a function of cell density ($OD_{600}$). Strains express *HXK1*, *HXK2*, or *E. coli glk* (EcGlk) from the constitutive pTDH3 promoter in an *hxk1*Δ *hxk2*Δ background. One biological replicate is shown.

**(B)** Doubling times determined from $OD_{600}$ growth measurements during the glucose depletion assay in **(A)**. One biological replicate is shown.

**(C)** Glucose consumption flux calculated from the glucose depletion measurements in **(A)** and the doubling times in **(B)**. One biological replicate is shown.

Cells were grown in media containing 10 mM glucose.

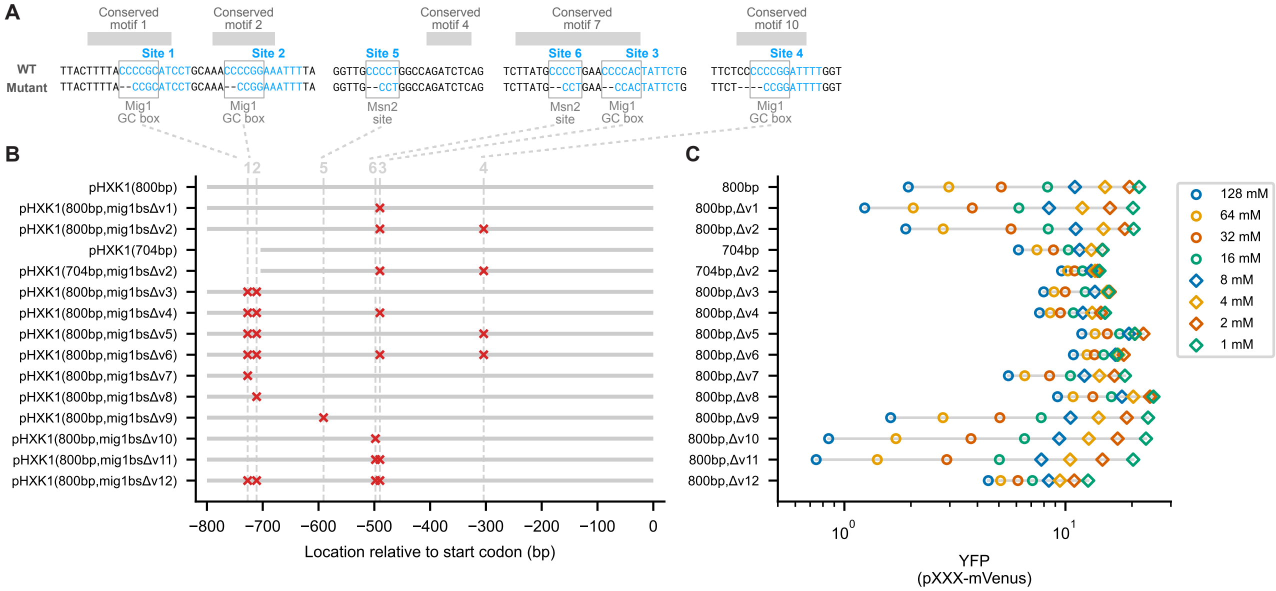


**Figure S9. Mapping Mig1 binding sites in the *HXK1* promoter, related to Figure 6.**
**(A)** Conserved sequence motifs identified by alignment of *HXK1* promoters from *Saccharomyces* species. Putative Mig1 and Msn2 binding sites within conserved regions are shown together with the nucleotide deletions introduced to disrupt each site.
**(B)** Schematic of the *HXK1* promoter variants used to map regulatory elements. Gray bars indicate promoter fragments, and red crosses indicate disrupted Mig1 or Msn2 binding sites.
**(C)** Steady-state activity of the *HXK1* promoter variants across a glucose titration. Each point represents the mean fluorescence averaged across at least two biological replicates.

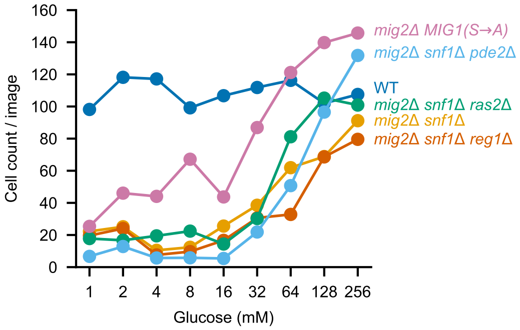


**Figure S10. Cell counts of upstream regulatory gene mutant strains, related to Figure 7.**
Average cell counts per image measured during fluorescence microscopy for the strains analyzed in Figure 7B–D across the glucose titration. Reduced cell counts at low glucose indicate growth defects in the corresponding mutant strains under these conditions. Points represent the mean of three biological replicates.

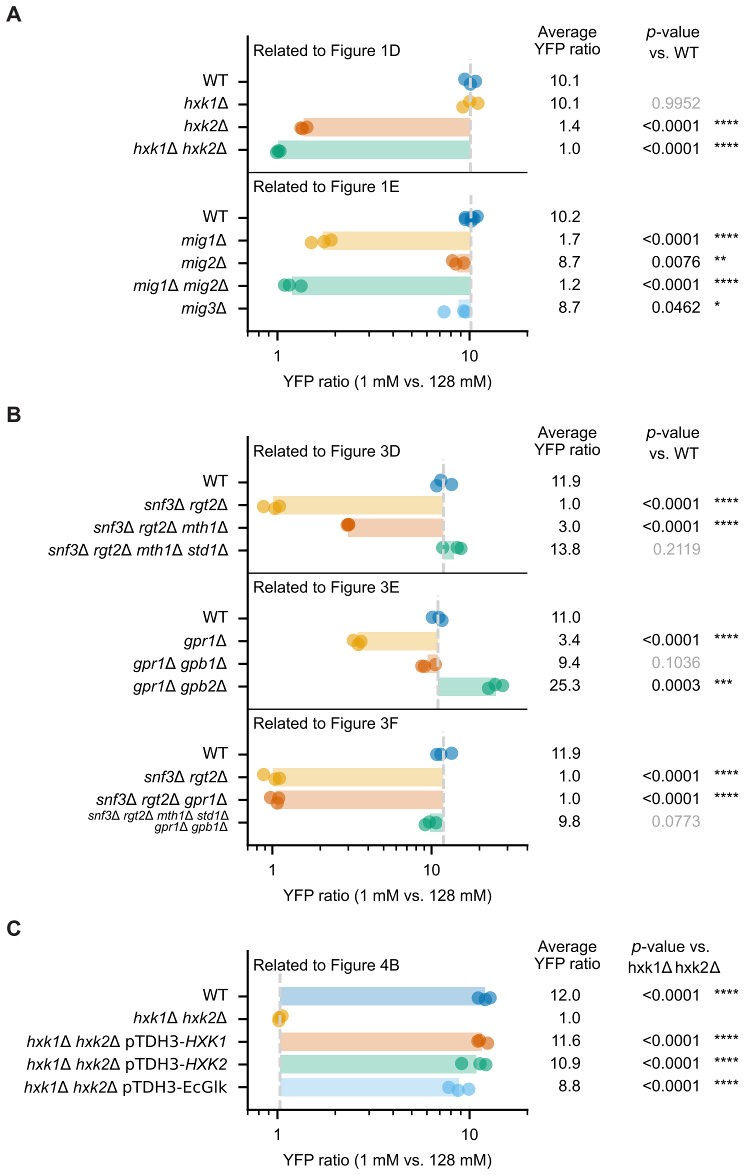


**Figure S11. Statistical analysis of glucose repression phenotypes, related to Figures 1, 3, and 4.**
**(A–C)** Quantification of glucose repression for the strains shown in Figures 1, 3, and 4, respectively. The x-axis shows the ratio of pHXK1-mVenus fluorescence measured at 1 mM versus 128 mM glucose. Each dot represents one biological replicate. The dashed line indicates the mean YFP ratio of the corresponding reference strain, and the colored bar indicates the fold change relative to the reference. Mean YFP ratios and $p$ values are listed on the right. Statistical significance was determined using two-sided independent two-sample Student's $t$-tests. $* \; p < 0.05$, $** \; p < 0.01$, $*** \; p < 0.001$, $**** \; p < 0.0001$.

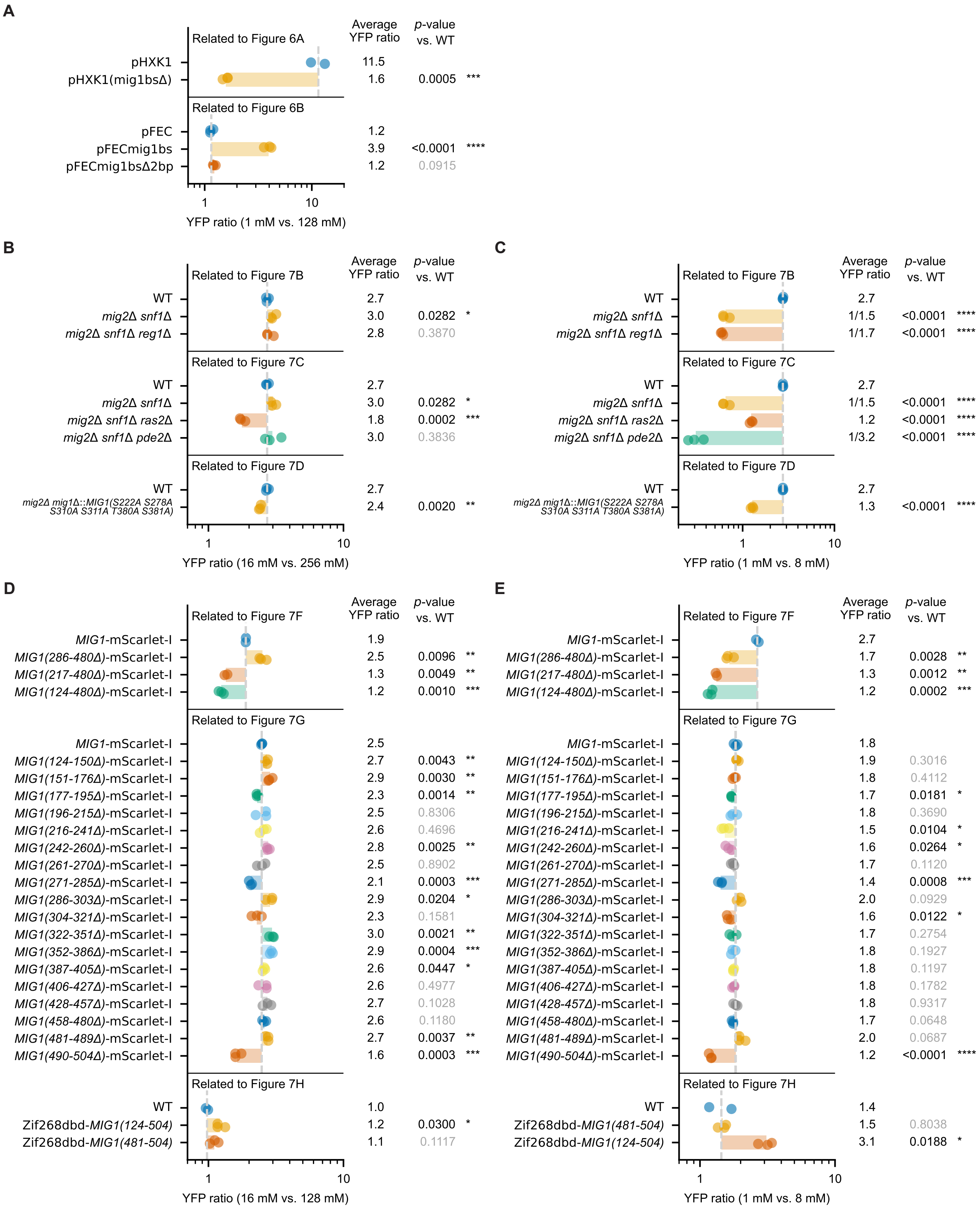


**Figure S12. Statistical analysis of glucose response phenotypes, related to Figures 6 and 7.**
**(A–E)** Quantification of glucose response for the strains shown in Figures 6 and 7. The x-axis shows the ratio of pFECmig1bs-mVenus or pZif268bs3-mVenus fluorescence measured in the full glucose range (1 mM versus 128 mM; **A**), the high-glucose range (16 mM versus 256 or 128 mM; **B**, **D**), or the low-glucose range (1 mM versus 8 mM; **C**, **E**). Each dot represents one biological replicate. The dashed line indicates the mean YFP ratio of the corresponding reference strain, and the colored bar indicates the fold change relative to the reference. Mean YFP ratios and $p$ values are listed on the right. Statistical significance was determined using two-sided independent two-sample Student's $t$-tests. $* \, p < 0.05$, $** \, p < 0.01$, $*** \, p < 0.001$, and $**** \, p < 0.0001$.
